# A Protein Language Model Reveals Organellar Ca^2+^ ATPases at Neuronal Synapses

**DOI:** 10.64898/2026.03.02.709014

**Authors:** Valentina Villani, Silvia Turchetto, Lars Boye Brandt, Markus Ørnsvig Christensen, Erika Uddström, Sara Derosa, Esben Lorentzen, Chao Sun

## Abstract

The synapse between neurons hosts the protein machinery for information transfer and storage in the brain. Its local proteome, however, contains many proteins with unclear roles for synaptic function. To glean hypotheses from this local proteome, we developed Synapse Gigamapper (SyGi), a protein language model for predicting protein localization at synapses. SyGi identified 152 amino-acid motifs that are indicative of localization at excitatory or inhibitory synapses and revealed >100 candidate constituents from key cellular pathways. Among these candidates is the endoplasmic reticulum (ER)-bound ATPase for cytosolic Ca^2+^ clearance, SERCA. We found clusters of SERCA copies at excitatory synapses without ER. Rather, SERCA is colocalized with compartment-specific organelles at synapses— the spine apparatus and synaptic vesicles. Taken together, SyGi is useful for exposing hidden components of neuronal synapses.

## Introduction

The synaptic contacts between neurons hosts the protein machinery for releasing and detecting neurotransmitters, enabling information transfer and storage in the brain. Their molecular content has been mapped by many experimental and expert-curated datasets (*1–4*). In recent studies, mass spectrometry routinely identifies over 2000 proteins in synaptic compartments dissociated from the mammalian brain (‘synaptosomes’) (*1*, *3–10*), with many identifications falling into the abundance range of false detections and contaminants. This technical limit may mask low-abundance proteins that *e.g.* exist in a subset of synapses— a potential driver of synapse heterogeneity (*11–13*). So far, expert-curated databases such as the Synaptic Gene Ontologies (SynGO) database only contain a subset of the protein identifications from proteomics (*4*)— many remain unaccounted for (Fig. 1A). This fraction of unassigned proteins remains up to ∼45% even for the proteomes of sorted specific synapse types (*1*), raising the possibility that many unassigned proteins are actual constituents of the synapse and carry out functions locally.

**Fig. 1.**
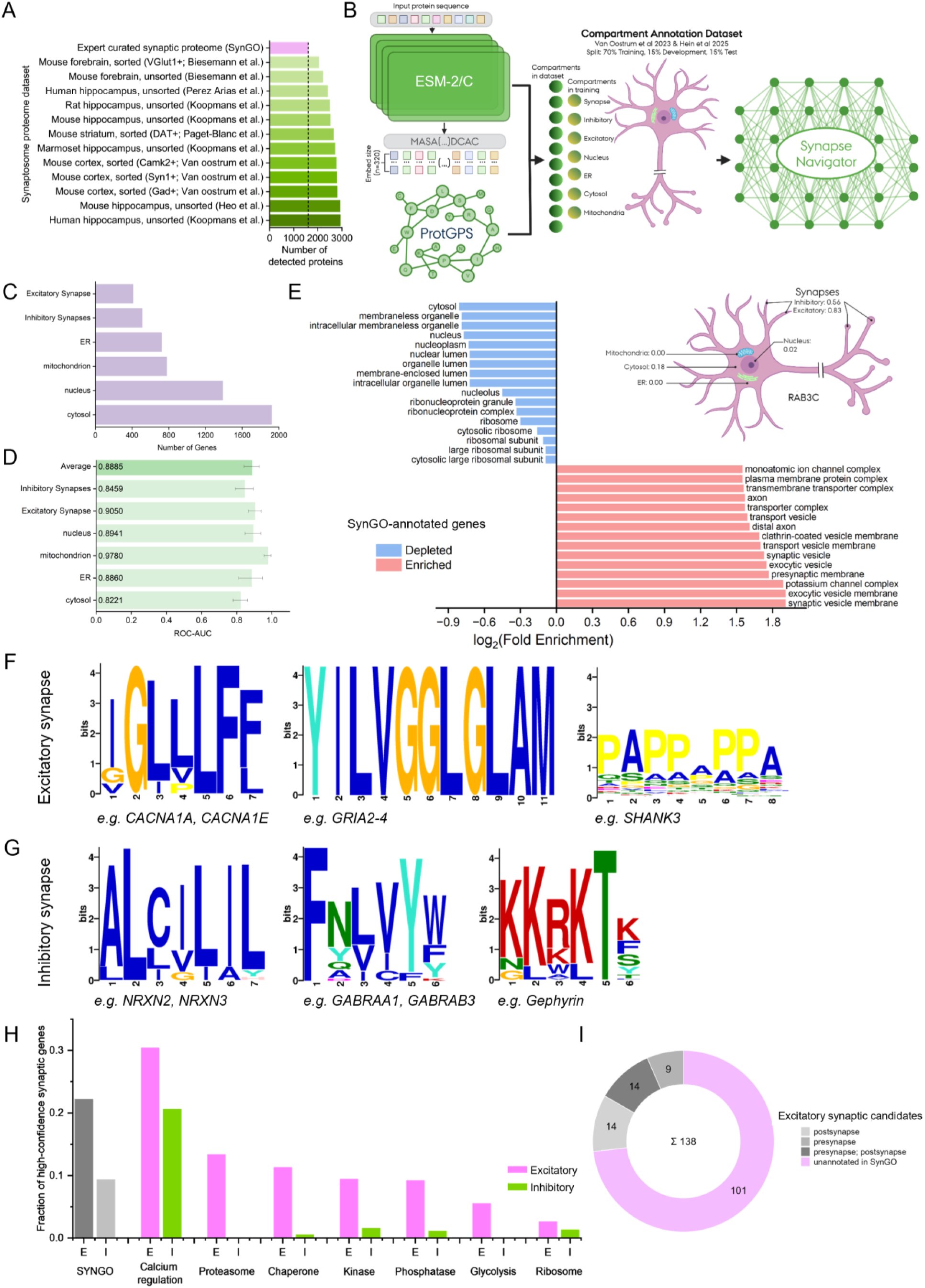
Synapse Gigamapper predicts protein localization in excitatory and inhibitory synapse types. (A) Bar graph showing that a large fraction of mass spectrometry-based protein identifications in unsorted or sorted synaptosomes is not yet accounted for in expert-curated synaptic protein databases such as SynGO (*1*, *4*, *6*, *7*, *9*, *10*). (B) Scheme illustrating the workflow of Synapse Gigamapper for predicting protein localization at excitatory and inhibitory synapses as well as four other essential cellular compartments. (C) Bar graph showing the number of genes that are annotated for each pre-defined subcellular compartment (see also Supplemental Data S1). (D) Bar graph showing the ROC-AUC values of SyGi performance in each pre-defined subcellular compartment. (E) Bar graph showing the Gene Ontology analysis of SynGO-annotated Genes that are significantly enriched or depleted by SyGi predictions (Panther Gene Ontology over-representation test against the complete SynGO gene list- cellular components; FDR < 0.05). Inset shows an exemplary synaptic gene Rab3C’s confidence score across all six subcellular compartments in neurons. (F&G) Logo plots of exemplar significantly overrepresented amino-acid motifs associated with localization at excitatory (F) and inhibitory (G) synapses according to SyGi. (H) Bar graphs showing the fraction of high-confidence (arbitrary cut-off ≥ 0.50) genes shortlisted by SyGi for excitatory (E; magenta) and inhibitory (I; green) synapse localization among seven curated functional protein families as well as SynGO-annotated genes (grey). (I) Pie chart showing that 101 out of 138 high-confidence genes in key cellular pathways in H predicted by SyGi for the excitatory synapse (light magenta) have not been annotated by SynGO as synaptic constituents.

This idea has accrued mounting evidence. For example, many ribosomal and proteasomal proteins have been identified in the synaptic proteome (*14*, *15*). Their expression and function at synapses allow protein synthesis and degradation to take place locally (*16–18*), enabling on-demand synaptic modification during memory formation (*19–22*). Recent studies further suggest that ribosomes and proteasomes also exhibit subcellular-specific features near synapses (*14*, *15*, *23*, *24*), raising possibilities of potential hidden proteins and protein function at synapses (*25*, *26*). To glean hypotheses from protein identifications in synaptic compartments at high throughput, we developed a transformer-based protein language model that can predict subcellular protein localization to excitatory and inhibitory synapse types based on amino-acid sequences, called Synapse Gigamapper (SyGi). Using SyGi, we identified 152 amino-acid motifs that are significantly enriched in excitatory and inhibitory synapse types, many of which are known localization signals for synaptic proteins. We further shortlisted protein candidates from key cell biological pathways for potential synapse function, leading to the discovery of clustered sarco/endoplasmic reticulum calcium ATPase (SERCA) expression at excitatory synapses. As a canonical component of the endoplasmic reticulum (ER), SERCA was found to localize to the spine apparatus— a specialized ER extension in the postsynaptic compartment— as well as >50% of synaptic vesicles in the presynaptic compartment. Overall, the protein language model here will be useful for discovering hidden constituents of neuronal synapses.

## Results

To develop a high-throughput method for selecting protein candidates that may localize to different synapse types, we developed SyGi, an evolutionary scale modelling (ESM)-based protein language model trained by published type-specific proteomic datasets of cortical excitatory and inhibitory synapses (*1*) as well as the subcellular proteomes of core cellular compartments: the nucleus, ER, mitochondria, and the cytosol (*27*) (Fig 1B; see also Supplemental Data S1). The training proteome dataset contains 6627 unique human UniProt IDs with experimentally validated subcellular localization annotations (Fig. 1C; see also Methods). Using this curated dataset, SyGi is built on the representational power of ESM-2/C embeddings and a published multi-task learning design (*28*), and can be customized on a standard desktop workstation equipped with a consumer-grade RAM and GPU within a couple of days (See Methods).

To benchmark the performance of SyGi, we calculated the ROC-AUC values of all six subcellular compartments (Fig. 1D). ROC-AUC values measured SyGi’s ability to rank true positives higher than true negatives (Fig. S1; see also Methods). Considering the class imbalance in gene number among the six compartments (Fig. 1C), ROC-AUC was used because it is robust to class imbalance and without arbitrary thresholds. Overall, across all compartments, SyGi achieved ROC-AUC values of > 0.8 (Fig. 1D & S1; averaged at 0.89 ± 0.03), on par with the state-of-the-art (*28*) (see also Methods). These results indicate that SyGi achieves high performance in ranking true positives over true negatives across six subcellular compartments.

How can SyGi be used for understanding protein localization at neuronal synapses? For each queried gene, SyGi outputs a confidence score of localization for each subcellular compartment (e.g. Rab3C; Fig. 1E inset). To evaluate whether SyGi can be used to shortlist synaptic proteins beyond the curated dataset, we used SyGi to predict the subcellular localization of an expert-curated gene list for synapses (SynGO) (*4*). A strict, arbitrary confidence cut-off of ≥ 0.5 retained 380 out of 1643 SynGO-annotated genes (23%), enriching Gene-Ontology (GO) terms for core synaptic components such as synaptic vesicles and ion-channel complexes while depleting those of housekeeping cellular components such as the cytosol and the nucleus (Fig. 1E). In addition, SyGi scores individual amino acids of a sequence based on their influences for the protein’s predicted localization (*28*). Combining this information with motif-based sequence analyses (*29*), we performed comprehensive motif discovery and enrichment amongst annotated genes of excitatory and inhibitory synapses (see Methods). SyGi identified 80 significantly enriched amino-acid motifs that are indicative of protein localization in excitatory synapses and 72 in inhibitory synapses, including known synaptic localization motifs such as proline-rich motifs in excitatory postsynaptic-density proteins (*e.g.* Shank; Fig. 1F right), as well as transmembrane motifs for receptors, ion channels and adhesion proteins (Fig. 1F left and middle & Fig. 1G left), poly-basic membrane-binding motifs for inhibitory postsynaptic density proteins such as gephyrin (Fig. 1G right), and potential PDZ-binding motifs (*e.g.* -SVVL; see Supplemental Data S3). 107 out of these 152 motifs are annotated in the Eurokaryotic Linear Motif (ELM) resource database (*30*). Overall, these results indicate that SyGi can be used to predict protein subcellular localization at synapses and identify potential localization motifs.

What affects the performance of SyGi in different subcellular compartments? We noticed that the cytosol compartment achieved the lowest ROC-AUC values (0.82) despite having the highest protein number for training (Fig. 1C&D), suggesting that high protein number during training is not sufficient for high performance. The inhibitory-synapse compartment, on the other hand, gave the lowest performance in reducing false negatives (Fig. S2B), suggesting highly conservative predictions. To explore whether the deep-learning model itself is a main limiting factor in SyGi’s performance at the inhibitory-synapse compartment, we also trained SyGi using a next-generation model containing 600 million parameters (ESM-C; versus 8 million in ESM-2) with weighted contributions from synaptic proteins to correct the class imbalance (see Methods). While ESM-C-based SyGi reduces false-negative predictions across all six compartments (Fig. S2B & S3), it elevates false-positive predictions for both mitochondrion and ER compartments (Fig. S2A). These results suggest that ESM-2-based SyGi is better suited for selecting high-confidence proteins while ESM-C-based SyGi is ideal if subsequent validations are inexpensive.

To test whether SyGi can accelerate the discovery of proteins from key cellular processes at excitatory and inhibitory synapses, we applied ESM-2-based SyGi prediction to seven curated functional protein families of essential cellular pathways, including calcium homeostasis, kinases, phosphatases, ribosomal proteins, proteasomal proteins, chaperones, and glycolysis (Fig. 1H). SyGi shortlisted 138 high-confidence protein candidates from the seven protein families above the strict confidence cut-off used in Fig. 1E, the majority of which are not yet annotated as synaptic proteins (Fig. 1I; see also Supplemental Data S2). A higher proportion of high-confidence candidates was found in excitatory synapses than in inhibitory synapses across all seven functional families (Figs. 1H&S4). For example, SyGi identified 49 key kinases for excitatory synaptic function (Fig. S5A) such as Eph receptor tyrosine kinases (e.g. EPHA2-5, EPHA7, EPHA8), Calcium/Calmodulin-dependent protein kinases (CAMK2, CAMK4), and mitogen-activated protein kinases (e.g. MAP2K6, MAPK8, MAP3K1); in comparison, only 7 predicted kinases for inhibitory synapses are found, 6 of which Eph receptor tyrosine kinases (Fig. S5B). In addition, SyGi shortlisted 25 phosphatases for predicted localization at excitatory synapses including e.g. protein tyrosine phosphatase receptors (PTPRs). 22 chaperones including *e.g.* HSP70, HSP90, and CCTs, 6 proteasomal proteins, 2 ribosome subunits (RPLP2 and RPS21), and 3 glycolytic enzymes were also predicted for excitatory synapses (Fig. S5A). For example, RPLP2 was previously identified as a ribosomal subunit that exchanges into existing ribosomes near synapses (*14*). Overall, these predictions provide a plethora of protein candidates for investigating proteostasis and signaling at synapses.

Are there common features in the vocabulary of amino-acid sequences associated with synaptic localization? To reveal the biophysical properties of synapse-specific amino-acid motifs and their reusability among different genes, we analyzed the repertoire of significantly enriched motifs at excitatory and inhibitory synapses (Fig. 2). The identified 152 motifs are between 4-16 amino acids in width, with 7 motifs shared between excitatory and inhibitory synaptic proteins (Fig. 2A&B). A UMAP analysis of their biophysical properties revealed five common types including polar/regulatory, charged/signaling, strongly charged, aromatic-rich, and hydrophobic/transmembrane (TM)-like (Fig. S6). We further visualized the similarity among the motifs that are excitatory-specific, inhibitory-specific, and shared (Fig. 2C; see Methods), revealing an absence of motif clustering by synapse types (see also Fig. S7), although motif sharing appears to be more common among genes of inhibitory synapses than those of excitatory synapses (Fig. 2C). A small number of motifs are shared promiscuously amongst hundreds of genes, with the majority shared by < 3 genes in both excitatory and inhibitory synapses (Fig. 2D, 2E & S8A). We identified 7 motifs that appear to be generic (>200 genes), including *e.g.* XXXSXXE, AXXSX, and KXXXK (Fig. 2E; X denotes any amino acid). For the remaining 145 motifs, the top three are acidic docking motifs (EEXXXLX), SH3-binding motifs (KXXPXXP), and nuclear-export-signal (NES) motifs (LXYXXL). Top synaptic hub genes (*e.g.* Gria, Nrxn, Bsn, Pclo, Shank) and their common motifs are visualized in a bipartite network and a heatmap (Fig. S8). Overall, SyGi enabled a systematic analysis of the amino-acid sequence vocabulary among synaptic proteins.

**Fig. 2.**
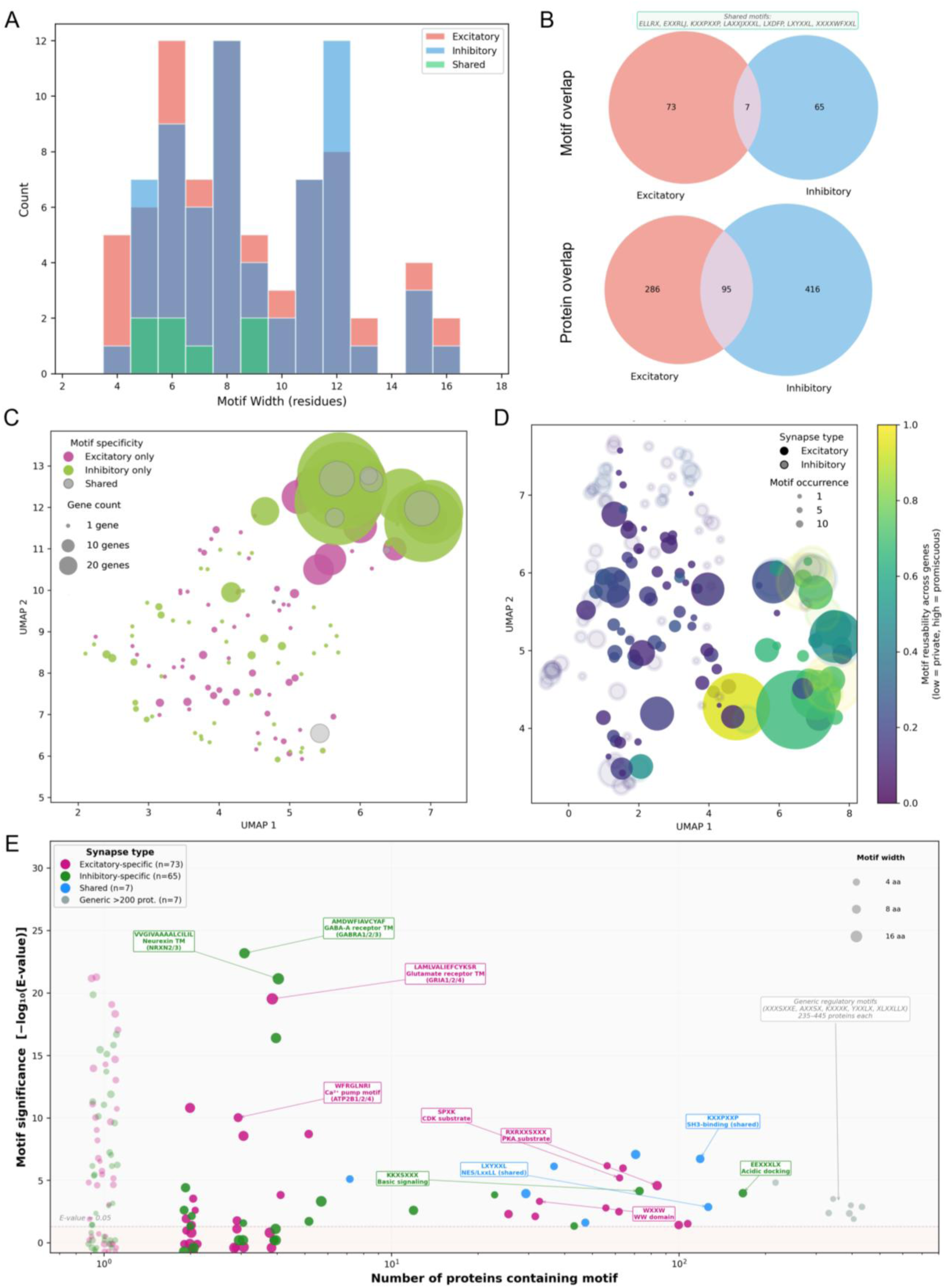
Synapse Gigamapper identifies 152 amino-acid motifs that are indicative of synaptic localization. (A) Bar graph showing the distribution of the widths (*i.e.* amino-acid number) of motifs that are significantly enriched in proteins from excitatory synapses (salmon), inhibitory synapses (cerulean), and both types (green). (B) Venn diagrams showing the number of shared motifs (in text box; X indicates any amino acid) and proteins that carry significantly enriched motifs. (C) UMAP plot showing the absence of clustering among motifs specific to excitatory (magenta) and inhibitory (green) synapses or shared among both (grey). Circle size indicates the number of genes that contain the motif. (D) UMAP plot showing the reusability and occurrence of motifs among genes of excitatory (filled circles) and inhibitory (hollow circles) synapses, where the circle size represents the total occurrences for each motif and the colors denote the reusability of a motif among genes (*i.e.* private vs promiscuous) calculated by Shannon entropy normalized by the number of genes (see Methods). (E) Scatter plot of 152 synaptic amino-acid motifs identified by SyGi, their specificity to synapse types (color), motif width (circle size), and exemplar genes and annotated motif functions (text boxes). Motifs found in isolated genes were denoted by circles with faded colors.

How can SyGi be used to generate hypotheses for synapse biology? For example, we noticed that the highest fractions of predicted genes for both excitatory and inhibitory synapses are found in the functional group of calcium homeostasis (Fig. 1H; >30% to excitatory synapses and >20% to inhibitory synapses). GO analyses of these candidates in excitatory synapses highlight the enrichment of genes involved in active calcium transport across ER and plasma membranes (PM). Consistently, a motif enrichment analysis following SyGi also identified a significantly enriched excitatory synaptic motif for calcium pumps (WFRGLNRI; Fig. 2E). These predictions warrant an experimental investigation of ER- and PM-bound calcium ATPases at excitatory synapses.

To investigate the synaptic presence of calcium ATPases, we visualized both the plasma-membrane calcium ATPase (PMCA) and the ER-bound calcium ATPase (SERCA2, the predominant brain isoform) via confocal microscopy of cultured, primary rat hippocampal neurons using validated antibodies (Fig S9)(*31*). Using vGluT1 as a reference marker for excitatory synapses, we found that PMCA appears to coat the neuronal surface, even around the vGluT1 puncta (Fig. 3B&D). We expected SERCA to distribute diffusely within the neuronal dendrites, considering the pervasive presence of ER along neuronal projections (*32*, *33*). However, we observed punctate SERCA distribution in dendrites that colocalized with vGluT (Fig. 3C&D), indicating a clustering of SERCA at excitatory synapses. Indeed, a significantly higher enrichment of SERCA at synapses versus PMCA was found (Fig. 3D). The divergent distribution patterns of the two major calcium ATPases near synapses were visualized in Fig. 3E, highlighting the punctate distribution of SERCA that decorates the dendritic branches (Fig. 3E inset). Using DNA PAINT-based single-molecule localization microscopy (*15*, *34*), we resolved the localization of individual PMCA and SERCA copies along the dendrites and at synapses (see Methods). DNA PAINT revealed clusters of SERCA copies at synapses (Fig. 3F), creating hotspots of SERCA localization on top of a pervasive distribution of PMCA (Fig. 3G). Despite the higher dendritic densities of detected PMCA copies (Fig. 3G; see Methods for DNA PAINT-based copy-number analyses), we estimated on average more SERCA copies per synapse (32.8±28.3 SERCA copies versus 19.5±25.0 PMCA copies) due to the synaptic clustering of SERCA (Fig. 3H). These results revealed clustering of SERCA at excitatory synapses. But how does ER cluster SERCA copies at synapses?

**Fig. 3.**
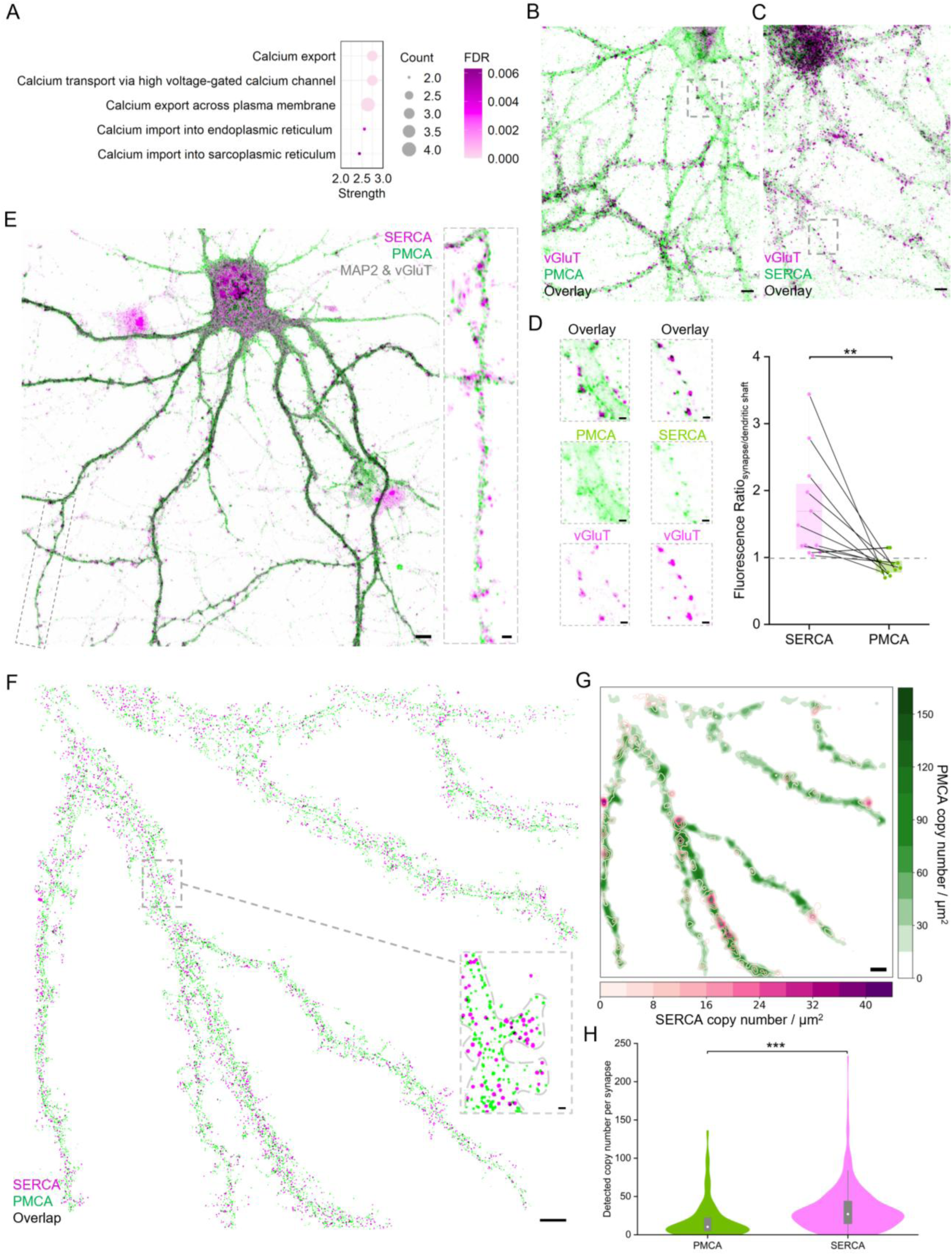
SERCA is clustered at excitatory synapses while PMCA exhibits diffuse presence along neuronal dendrites. (A) Gene ontology analysis (DAVID-overrepresented biological processes) of Top-5 enriched terms among calcium regulators predicted to localize at excitatory synapses. (B) & (C) Confocal micrographs of neuronal PMCA (green; B) and SERCA (green; C) in addition to vGluT1 (magenta) in cultured neurons (soma at the top). Scale bars: 5 μm. (D) Column scatter plots showing SERCA is enriched at vGluT1-positive synapses versus the dendritic shaft at a ratio that is significantly higher than PMCA. (Averaged across synapses from 15 neurons; paired two-sample T-test, with p**<0.01). Left inset shows exemplary synaptic localization of PMCA and SERCA from the regions in the dashed boxes from B&C. Scale bars: 1 μm. (E) Confocal micrograph of neuronal PMCA (green) and SERCA (magenta) along dendrites and at synapses (Map2 & vGluT; grey). Scale bar: 4 μm. Inset highlights the puncta expression pattern of SERCA along a dendrite branch versus the diffuse presence of PMCA. Scale bar: 1 μm. (F) DNA PAINT-based single-molecule localization micrograph of neuronal PMCA and SERCA copies (green and magenta, respectively, with overlap in black) along dendrites & at synapses. Scale bar: 2 μm. Inset shows exemplar dendritic spines with resolved SERCA and PMCA copies. Scale bar: 200 nm. (G) Overlay between the heatmap for PMCA copy distribution and the contour map for SERCA distribution along the neuronal dendrite. The color scales indicate the detected copy-number densities of SERCA (magenta) and PMCA (green) per μm^2^ dendritic area (MAP2 mask). Scale bar: 5 μm. (H) Violin plot showing a significantly higher amount of detected SERCA copies than PMCA copies per vGluT-positive synapse in F (647 synapses from 6 neurons; two-sample T-test; p***<0.001; white dots denote median values).

To address whether ER might host SERCA clusters at synapses, we visualized the ER and SERCA in neuronal projections via immunofluorescence confocal microscopy (Fig. 4A). Using calnexin as an ER marker (cyan; Fig. 4A), we found that ER is prevalent along neuronal processes but rarely enters dendritic spines, consistent with previous studies (*32*, *33*, *35*). While many dendritic SERCA colocalize with calnexin, a subpopulation of SERCA signals was colocalized with vGluT1 puncta in the absence of calnexin (indicated by grey arrows; Fig. 4A inset). In fact, the SERCA intensity at vGluT1-positive areas was not significantly different from that at calnexin-positive areas (Fig. 4B), suggesting an alternative host for synaptic SERCA clusters.

**Fig. 4.**
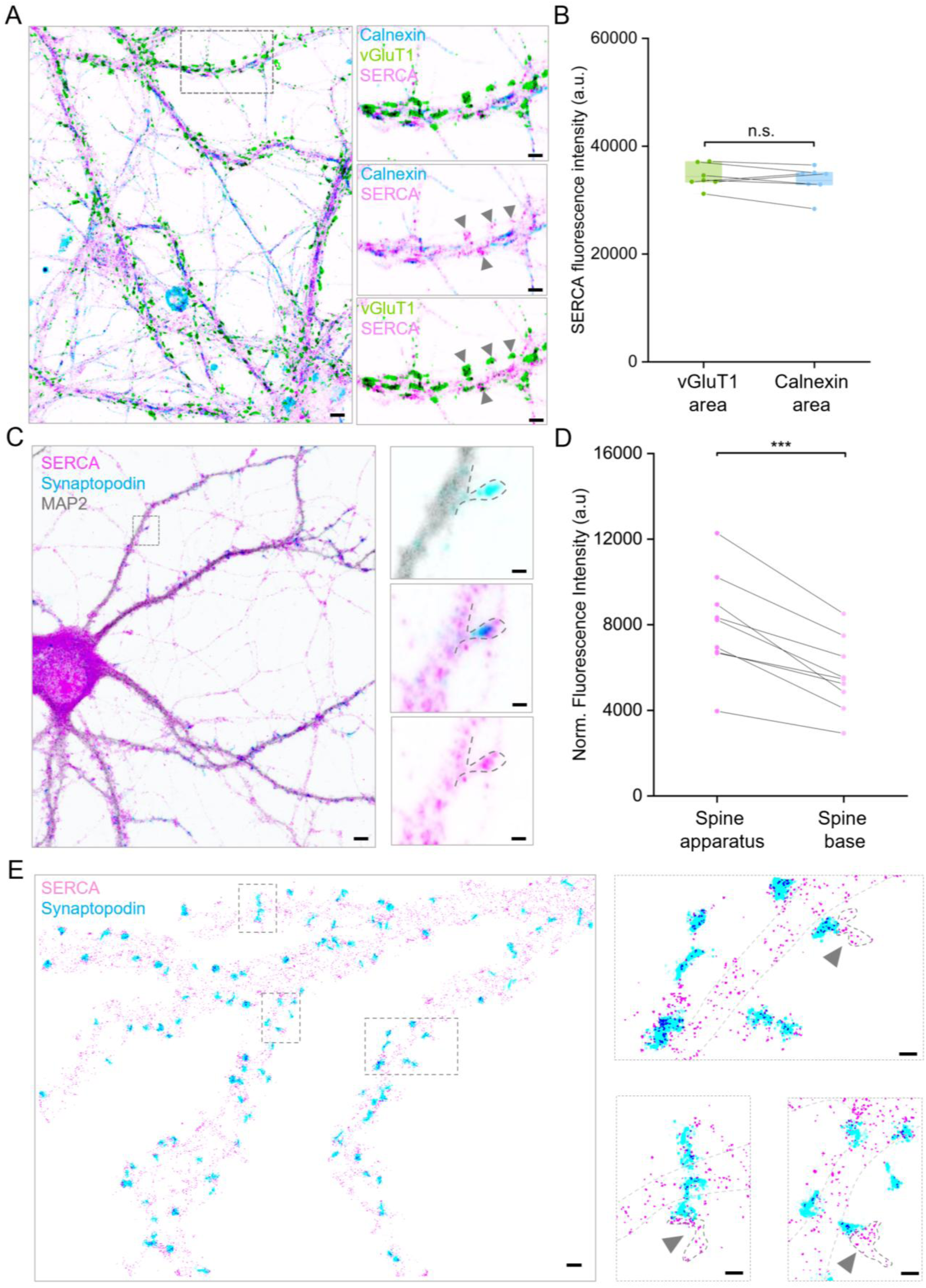
Synaptic SERCA clusters are hosted by the spine apparatus in the postsynaptic compartment. (A) Confocal micrograph showing the distribution of ER (anti-calnexin; cyan), excitatory synapses (anti-vGluT1; green), and SERCA (magenta) in neuronal processes, with part of the cell soma in the bottom-right corner). Scale bar: 1 μm. Right inset shows a cluster of excitatory synapses containing calnexin-negative SERCA fluorescence (indicated by grey arrows) in addition to SERCA signal that colocalized with Calnexin (blue for overlap between SERCA and ER). Scale bars: 1 μm. (B) Box plot showing comparable levels of SERCA fluorescence (normalized by area) at vGluT1-positive areas versus Calnexin-positive areas in neuronal dendrites (Paired two-sample T test- p>0.05; averaged across synapses from 15 neurons from 3 neuronal preparations per group). (C) Confocal micrograph showing the co-localization of synaptic SERCA clusters (magenta) with the spine apparatus (anti-synaptopodin; cyan) in cultured neurons, with overlay in blue. Scale bar: 2 μm. Inset shows a dendritic spine containing the spine apparatus (cyan) and a cluster of SERCA (magenta), with MAP2 in grey. Scale bars: 0.5 μm. (D) Box plot showing significantly higher levels of SERCA fluorescence (normalized by area) in synaptopodin-positive areas versus the spine base (two-sample T test- p***< 0.001; averaged across synapses from 15 neurons from 3 neuronal preparations per group). (E) DNA PAINT-based single molecule localization micrograph of SERCA and the spine apparatus in cultured neurons. Scale bar: 1 μm. Right inset shows exemplary spines (indicated by grey arrows), where some synaptic SERCA copies (encircled by dark grey dashed outlines) are neither in the dendritic shaft (outlined by lighter grey dashed lines) nor in the spine apparatus (cyan). Scale bars: 1 μm.

To elucidate the local host of synaptic SERCA clusters, we visualized the specialized ER extension that resides in a subpopulation of dendritic spines, the spine apparatus (*36*). It has been long speculated to participate in local calcium regulation (*36*) during plasticity (*37–39*), and its stereotypical stacks of smooth ER cisterns appears ideal for gathering clusters of SERCA copies (*40*). Indeed, in Fig. 4C, we observed colocalization of SERCA and synaptopodin (reference marker for the spine apparatus) via immunofluorescence confocal microscopy. A significantly higher level of SERCA was found present at the spine apparatus versus the spine base. In contrast, the Ryanodine receptor (RyR), an ER-bound calcium channel, is present albeit not clustered at the spine apparatus (Fig.S10B-D). Compared to SERCA, RyR exhibited a significantly lower fraction of dendritic copies at the spine apparatus (Fig. S10E), consistent with its low confidence score for synaptic localization by SyGi (see Supplemental Data S2) as well as previous studies (*35*). At single-protein resolution, DNA PAINT resolved the SERCA copies at the spine apparatus (Fig. 4E), revealing an overlap between the hotspots of the spine apparatus and the contours of dendritic SERCA copies (Fig. S10A). These observations revealed that the spine apparatus, a specialized ER extension into dendritic spines, hosts SERCA clusters in the postsynaptic compartment.

While many SERCA copies are colocalized with synaptopodin, the single-protein resolution of DNA PAINT also revealed synaptic SERCA copies adjacent to synaptopodin and, yet, outside the dendrite (Fig. 4E right inset; indicated by dark gray dashed outlines), suggesting a presynaptic presence of SERCA. To elucidate the compartment-specific expression of SERCA in pre- and postsynaptic compartments, we visualized the synaptic expression of SERCA together with pre-and postsynaptic markers (vGluT1 and PSD95, respectively) via 4X expansion microscopy of fixed mouse brain slices (Fig. 5A). The ∼4X expansion was validated using the sizes of nuclei stained by DAPI (Fig. S11). Consistent with *in vitro* (Fig.3), SERCA clusters were colocalized with both vGluT (overlap in yellow; Fig. 5B) and PSD95 (overlap in red; Fig. 5B) at excitatory synapses of the mouse cortex. Intriguingly, a significantly higher level of SERCA was detected in the presynaptic compartment than in the postsynaptic compartment (Fig. 5C). If the spine apparatus is a host of synaptic SERCA clusters in the absence of ER, why is their presynaptic level higher than the postsynaptic?

**Fig. 5.**
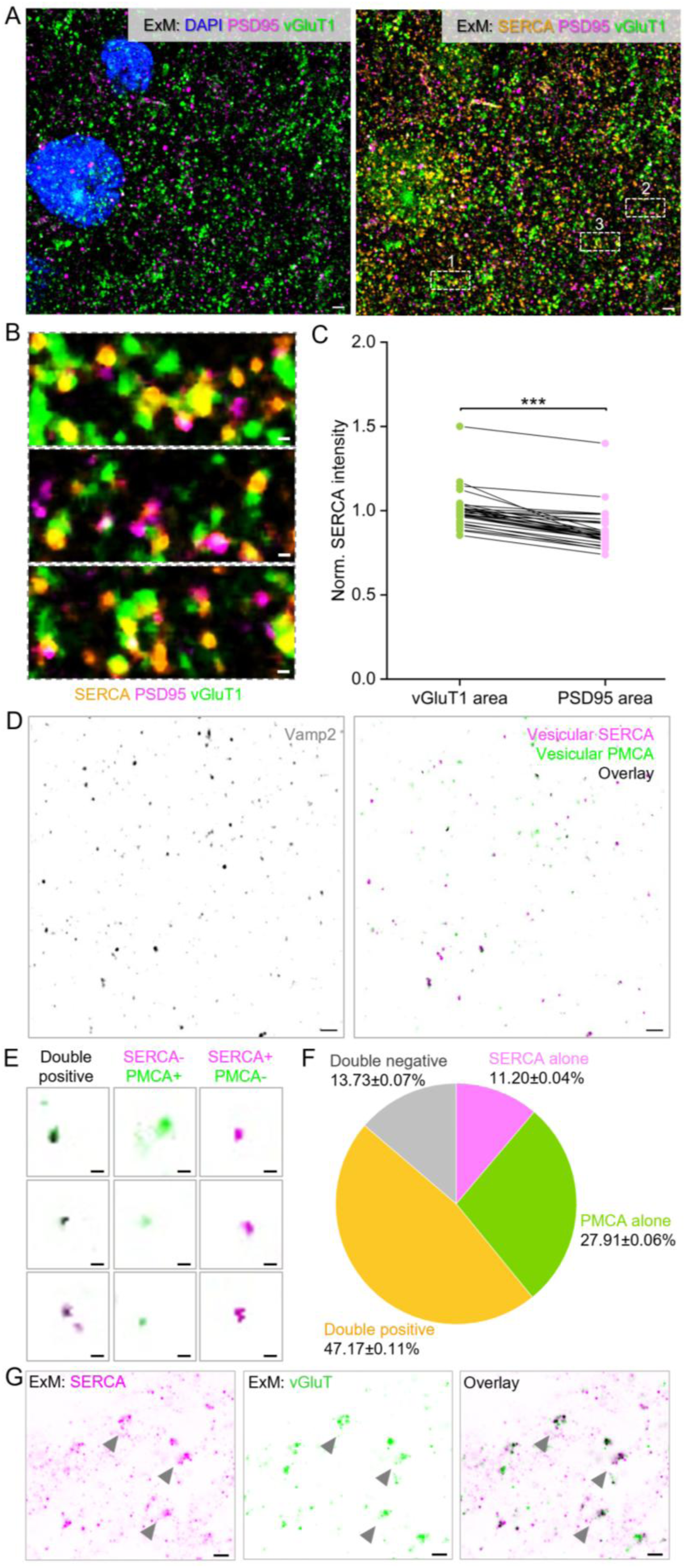
Presynaptic SERCA clusters are localized at synaptic vesicles. (A) 4X Expansion Microscopy of SERCA (orange), PSD95 (magenta), vGluT1 (green), and DAPI (blue) in the cortical area of a fixed mouse brain slice. Scale bars: 4 μm. (B) Magnified views of neuronal synapses (indicated by numbered boxes in the right image of A) containing SERCA signals (orange). Overlap between PSD95 (magenta) and SERCA is red. Overlap between vGluT1 (green) and SERCA is yellow. Scale bars: 1 μm. (C) Column scatter plots showing a significantly higher level of SERCA expression (normalized by area) in vGluT1-positive regions than in PSD95-positive regions (p***<0.001; Paired two-sample T-tests; 32 fields-of-view from 3 slice preparations; normalized against average values in vGluT1 areas). (D) Confocal micrographs of SERCA and PMCA signals (magenta and green, respectively, with overlay in black; right) masked by Vamp2 signals (grey; left) using purified synaptic vesicles plated on glass. Scale bars: 2 μm. (E) Exemplar diffraction-limited micrographs of Vamp2-positive puncta that are positive for both SERCA & PMCA (double-positive) or only one of them. Scale bars: 300 nm. (F) Pie chart showing the fractions of Vamp2-positive puncta from plated purified synaptic vesicles that carry SERCA and/or PMCA averaged across 6 fields of view from 3 preparations. (G) 4X expansion microscopy showing *in situ* colocalization of SERCA (magenta) with vGluT (green) at the excitatory synapses of cultured neurons. Scale bars: 2 μm.

We hypothesized that SERCA may also form clusters at the presynaptic compartment independent of ER. Previous studies have proposed the idea that calcium transport into synaptic vesicles is important for presynaptic Ca^2+^ clearance during activity (*41*). Consistently, both SERCA and PMCA have been identified in purified synaptic vesicles, albeit at depleted levels in comparison to input (*42*, *43*). To directly test whether SERCA and PMCA may be present at synaptic vesicles, we visualized SERCA and PMCA in *ex vivo* synaptic vesicles purified from the mouse brain in a previous study (gift from Rubinstein lab; see Methods) (*43*). Confocal microscopy of the purified synaptic vesicles plated on glass revealed that 86% of Vamp2-positive puncta possess fluorescence signals for either SERCA or PMCA (Fig. 5D-F), with 47% puncta exhibiting both SERCA and PMCA signals (Fig. 5E&F; double positive). As negative controls, postsynaptic proteins PSD95 and Homer1 were both found undetected on Vamp2-positive synaptic vesicles following immunofluorescence staining (Fig. S12; vs. synaptophysin as positive control). Furthermore, to validate the vesicular presence of synaptic SERCA *in situ*, we performed 4X expansion microscopy of SERCA and vGluT1 in cultured neurons, revealing a tight colocalization between SERCA puncta and vGluT1 puncta (Fig. 5G). Finally, the immunofluorescence of vesicular SERCA was observed without permeabilization prior to staining by an antibody that targets the cytoplasmic epitopes of SERCA (Fig. S12D&E), suggesting that its cytoplasmic domains remain in the cytosol. Overall, these results demonstrated that a canonical ER-bound calcium ATPase is localized to synaptic vesicles in the presynaptic compartment and the spine apparatus in the postsynaptic compartment, leading to its clustering at neuronal synapses via compartment-specific mechanisms.

## Discussion

To enable parallel information processing among synapses, the neuron has decentralized a variety of cell-biological machinery to synapses, allowing complex subcellular biology in the local milieu. To uncover these subcellular processes, artificial intelligence-based high-throughput approaches can accelerate biological discovery by selecting candidate genes and pathways among thousands of protein identifications at neuronal synapses. Here we used a protein language model to shortlist protein candidates that may localize to excitatory and inhibitory synapses. SyGi not only identified hundreds of amino-acid motifs that correlate with synaptic localization but also shortlisted many candidate proteins from key cell-biology pathways as potential synaptic constituents. Its hypothesis-generating power is evident in the discovery of compartment-specific mechanisms for hosting intracellular calcium ATPases at excitatory synapses.

SyGi can be used for high-throughput *in silico* screening of hidden genes at neuronal synapses. Its performance appears to be more limited by the quality and quantity of biological data rather than by the power of protein language models. For example, when a pre-defined subcellular compartment does not possess a highly compartmentalized, local proteome (*e.g.* the cytosol, an intermediate station for many proteins destined for other compartments), SyGi tends to underperform. The low performance for the inhibitory-synapse compartment may also be affected by this factor, considering that inhibitory synapses are largely shaft synapses with a much less compartmentalized postsynaptic milieu compared to excitatory synapses (*44*). Furthermore, when a pre-defined subcellular compartment contains a limited number of proteins for model training (*i.e.* the inhibitory-synapse compartment in Fig. 1C), SyGi’s performance may also be compromised. Likely for these reasons, we have not succeeded in including dopaminergic synapses or Golgi as viable compartments for SyGi.

ESM-2 based SyGi and ESM-C based SyGi provide complementary strengths depending on the cost of carrying false positives and false negatives in downstream experimental validations. ESM2 is ideal for generating high-confidence shortlists that will be tested in expensive or time-consuming follow-up experiments while ESM-C is ideal for ensuring comprehensive coverage before high-throughput experimental validations. Despite the conservative predictions of ESM-2 based SyGi, we generated >100 candidate proteins for the synapse, providing numerous testable targets. The identification of 152 amino-acid motifs also expands the repertoire of potential synaptic targeting cues, revealing the scope of potential primary sequences for organizing neurons— a cell type that relies on extreme compartmentalization to operate the complex neural network. In sum, protein language models such as SyGi have become relevant for accelerating experimental neurobiology.

SyGi can provide fresh perspectives on huge volumes of experimental data that have accumulated. For example, the proteome of purified synaptic vesicles has been investigated extensively in literature (*42*, *45*), and both SERCA and PMCA were identified, albeit at depleted levels compared to input (*42*, *43*). Their depleted abundance in synaptic vesicles is unsurprising considering that SERCA and PMCA are primarily associated with the ER and the plasma membrane. In practice, an absence of enrichment in subcellular proteomics often means that these low-abundance protein identifications will be discarded due to suspected false identification. For similar reasons, many low-abundance protein identifications from isolated synaptosome preparations have not been annotated in the proteome of an ‘averaged’ synapse (*2*, *3*). However, low-abundance proteins can be a major driver of synapse diversity, imparting a subset of synapses with unique properties (*12*, *13*, *46*). SyGi may provide a rapid, high-throughput approach to generate hypotheses and salvage these ignored targets from experimental data.

Calcium clearance, an essential housekeeping mechanism for mammalian cells, appears to be delegated to specialized organelles at neuronal synapses. It is known that calcium activity is highly compartmentalized at excitatory synapses (*47*). This compartmentalization sets a requirement for rapid, local, cytosolic calcium clearance during high-frequency synaptic transmission (*48*). The clustering of SERCA at the spine apparatus meets this requirement, providing a structural basis for the role of synapotodin in Ca^2+^ storage and structural plasticity of dendritic spines (*38*); its positioning at spines may also gauge the thresholds of long-range Ca^2+^ signal transduction through dendritic ER (*35*). Similarly, the presence of SERCA on synaptic vesicles also provides a close-up sink for calcium at the presynaptic active zone, considering that the axonal ER is often detected at a distance from the active zone in the crowded presynaptic bouton (*33*). Speculatively, vesicular uptake of cytosolic calcium via SERCA and PMCA may provide negative feedback for calcium-driven vesicle fusion, shuttling presynaptic calcium back to the extracellular space in an activity-dependent manner. Finally, the detection of vesicular SERCA suggests that the ER may provide more than just lipids for synaptic vesicle assembly (*49*, *50*). These results reveal many intriguing leads for understanding the unique molecular logistics of neuronal synapses during information processing.

## Acknowledgments

We thank H.R Kilgore and R. Barzilay for input on protein language model training; L. Benedetti, T. A. Ryan, & J. Lippincott-Schwartz for input on ER imaging; J. Rubinstein and C. Coupland for the kind gift of purified synaptic vesicles; P. Nissen, E.M. Schuman, and L.Y. Wang for discussions; A. Banka for contributions to motif analyses; S. tom Dieck and L. N. tom Dieck for input on the spine apparatus; T. Birch for preparation of primary neuron cultures.

## Author contributions

Conceptualization: CS

Methodology: CS, EL, ST

Investigation: VV, ST, LB, MC, EU, SD, CS

Visualization: CS, EL, VV, ST, LB, MC, EU, SD

Funding acquisition: CS, EL, ST

Project administration: CS

Supervision: CS

Writing – original draft: CS, EL

Writing – review & editing: CS, EL, VV, ST, MC, EU, LB, SD

## Data, code, and materials availability

All data are available in the main text or the supplementary materials. Code, trained models, and analysis scripts are available at GitHub repository: https://github.com/Synaptic-Logistics-Lab/SynapseGigamapper

## Supplementary Materials

### Materials and Methods

#### Synapse Gigamapper (SyGi)

1. Model architecture: SyGi is a deep learning framework for predicting protein localization to cellular compartments with a specialized focus on two major types of neuronal synapses, including the cytosol, endoplasmic reticulum (ER), mitochondrion, nucleus, excitatory synapse, and inhibitory synapse. The model architecture consists of two components: (i) a protein language model encoder that generates contextualized sequence embeddings, and (ii) a multi-layer perceptron (MLP) classifier that predicts multi-label compartment assignments.

2. Protein language model encoder: SyGi uses ESM2-t6-8M embeddings (320 dimensions, averaged across residue positions) passed through a two-layer MLP classifier (512 hidden units, ReLU, BatchNorm, Dropout p=0.1) with sigmoid output for multi-label classification across six compartments (*28*). Total parameters: ∼8.4M. In parallel, we also built SyGi using ESM-C (EvolutionaryScale, 2024) with 600M parameters to test whether encoder capacity limits performance.

3. Training dataset: The dataset was constructed by integrating proteomic data from two complementary published resources: a global organelle profiling dataset (*27*) and a type-specific synaptic proteome dataset (*1*). For the synapse dataset, proteins enriched in cortical CamKII-positive and cortical GAD2-positive synaptosomes were selected to represent excitatory and inhibitory synapses, respectively. The protein identifications were mapped to their human orthologs using UniProt’s online ID-mapping tool, and only one-to-one matches were retained. The dataset comprised 7011 human proteins with experimentally validated subcellular localization annotations from UniProt (SwissProt and TrEMBL databases). Proteins were annotated for 15 distinct compartments based on Gene Ontology (GO) Cellular Component terms and curated localization data. 5% of the dataset (384 proteins) was excluded for final performance testing. The remaining 6627 proteins were used for training SyGi. We focused on six target compartments relevant to neuronal function:

I. Cytosol (n=1,806, 27.3%)
II. Endoplasmic Reticulum (n=680, 10.3%)
III. Mitochondrion (n=724, 10.9%)
IV. Nucleus (n=1,319, 19.9%)
V. Excitatory Synapse (n=413, 6.2%)
VI. Inhibitory Synapse (n=512, 7.7%)

Each protein was encoded as a binary vector indicating localization presence (1) or absence (0) across compartments, allowing multi-label annotations, as previously shown (*28*). Related categories were merged to simplify the structure: actin cytoskeleton, p-body, and cytosol were combined into cytosol, while ER and ERGIC were merged into endoplasmic reticulum (ER). The two datasets were merged, combining duplicate entries with different annotations into single multi-label entries to reflect potential dual localization. The dataset of 6627 proteins was split into training (70%, n=4,639), validation (15%, n=994), and test (15%, n=994) sets with stratification to maintain compartment frequency distributions across splits. Protein sequences ranged from 41 to 7,818 amino acids (mean: 638 aa, median: 472 aa), with sequences longer than 1,800 aa truncated during training. We note that class imbalance is a challenge inherent to the training dataset: Synapse-associated proteins represent only ∼14% of the total dataset (925/6,627), compared to 27% for cytosolic proteins. A 4-fold imbalance, combined with the small absolute number of synaptic proteins, poses a significant challenge for machine learning models, which tend to optimize overall accuracy by favoring abundant classes at the expense of rare ones. For this reason, ESM-C-based SyGi used 10X weight for inhibitory synapses and 8X weight for excitatory synapses.

4. Training Configuration: Training Configuration: SyGi was trained using PyTorch 2.0.0 and PyTorch Lightning 1.6.4 with the following hyperparameters: Adam optimizer (learning rate: 1×10⁻³), batch size: 10, maximum epochs: 30, learning rate scheduler: ReduceLROnPlateau (patience: 5 epochs, factor: 0.5), early stopping: patience of 10 epochs based on validation ROC-AUC, gradient clipping: max norm 1.0, maximum sequence length: 1,800 amino acids (longer sequences truncated), FP32 precision. Training was performed on a standard desktop workstation equipped with a consumer-grade GPU (NVIDIA GeForce RTX 4070) and RAM (128 GB), completing in ∼90 minutes (stopped at epoch 20 based on early stopping). The model checkpoint with the best ROC-AUC value (averaged across 6 compartments) was selected.

5. Model optimization: To address the poor recall for the inhibitory synapse compartment observed with the ESM-2 model (Fig. S2B), we explored whether a larger architecture combined with weighted loss could improve minority class detection. We tested a next-generation protein language model (ESM-C 600M, EvolutionaryScale 2024) with substantially larger capacity (600 million parameters, 1152-dim embeddings, 36 transformer layers). We implemented a weighted binary cross-entropy loss function with compartment-specific positive class weights. Weight hyperparameters were optimized through systematic grid search testing 14 different configurations (6 weight tuning variants, 2 focal loss variants, 1 two-stage approach, and 5 refinements). The optimal configuration used weights of 10× for excitatory synapses and 8× for inhibitory synapses, with standard weights for general compartments (2.67× cytosol, 8.75× ER, 8.15× mitochondrion, 4.02× nucleus). Training was performed on an NVIDIA A1000 80GB GPU.

Training configuration for ESM-C 600M:

– Batch size: 8 (reduced due to model size)
– Learning rate: 1×10⁻⁴ (reduced for stability)
– Mixed precision training (FP16)
– All other hyperparameters identical to ESM2-8M

6. Model Evaluation Metrics

Model performance was evaluated using multiple complementary metrics to account for class imbalance:

a. ROC-AUC: Receiver operating characteristic area under the curve, a measure of the model’s ability to discriminate between positive and negative classes across all classification thresholds (Fig. S1). Values range from 0.5 (random guessing) to 1.0 (perfect discrimination). A score of 1.0 is perfect; a score >0.8 is excellent; a score of >0.7 is acceptable; a score of 0.5 means no discrimination (*28*). ROC-AUC is threshold-independent and robust to class imbalance, making it ideal for comparing models.

b. Precision: True positives / (True positives + False Positives) (Fig. S2A)

c. Recall: True positives / (True positives + False Negatives) (Fig. S2B)

Evaluation protocol: All metrics were computed on the held-out test set (5% of total input, n=384 proteins) that was never seen during training or hyperparameter tuning. To avoid data leakage, no test set information was used for model selection or threshold optimization - all decisions were based solely on validation set performance. To include other standard multi-label classification metrics including precision and recall (Fig. S2), arbitrary thresholds of probability scores needed to be applied. Based on the Gene Ontology Analyses of SynGO-annotated genes (Fig. 1E), we defined predictions with a stringent arbitrary confidence score ≥ 0.50 as positive (1) and those < 0.50 as negative (0) in order to select high-confidence candidates for experimental validation.

Following model evaluation, additional analyses were performed to interpret predicted localization trends. Confidence-score distributions and ROC-AUC visualizations were generated using OriginPro 2025. For functional exploration, curated protein families (calcium regulators, kinases, ribosomal proteins, proteasomal proteins, phosphatases, glycolytic enzymes, and chaperones) were compiled from public databases and literature sources (see Supplemental Data S2). Predictions for these families were compared across compartments. Functional enrichment analyses were performed using STRING v12.0 for protein-protein interaction mapping and Gene Ontology (GO) annotation, complemented by the Panther GO Overrepresentation Test using the associated protein family dataset for independent statistical validation. Enrichment was assessed using Fisher’s Exact Test with false discovery rate (FDR) correction for multiple testing. STRING networks were visualized with the strength parameter (log10(observed/expected)) to indicate enrichment magnitude.

7. Sequence-Level Feature Analysis: SyGi calculates an attribution score for each amino acid in the protein sequence based on their contribution to the predicted subcellular localization, as previously reported (*28*). Based on these attribution scores, we performed XSTREME analyses (*29*), a motif discovery and enrichment framework within the MEME Suite, designed to identify short linear motifs in biological sequences across multiple permutations of sequence windows (the width of the analyzed sequence segment) and paddings (flanking amino acid residues) based on Expectation values (E-values; y-axis of Fig. 2E) of the MEME Suite. To validate the biological relevance of learned features, we analyzed amino-acid composition enrichment in synaptic vs. non-synaptic proteins (Fig. S2C). For each of the 20 standard amino acids, we calculated:

Enrichment ratio = Frequency in synapse proteins/Frequency in non-synapse proteins

Statistical significance was assessed using two-sample t-tests comparing amino acid frequencies between synapse-localized (n=925) and non-synapse proteins (n=5,702), with Bonferroni correction for multiple testing (α = 0.05/20 = 0.0025). Enrichment ratios >1.0 indicate amino acids overrepresented in synaptic proteins, while ratios <1.0 indicate depletion.

#### Cell culture

Dissociated rat primary hippocampal neuron cultures were prepared and maintained as previously described (*16*). In brief, rat hippocampal neurons from postnatal-day-1 rat pups of either sex (RRID:RGD_734476; strain Sprague-Dawley) were dissected and dissociated by incubating with L-cysteine-papain solution at 37°C before being plated onto 35-mm MatTek dishes (MATTEK, A Bico Company) coated with poly-D-lysine. Cultured neurons were incubated in Neurobasal A medium (Gibco) supplemented with B-27 (Invitrogen) and Glutamax (Gibco) at 37°C and 5% CO2 for 18 days before use. All procedures followed national animal care guidelines and the guidelines issued by Aarhus University and local authorities.

#### Western Blotting

Dissociated rat primary cortical neurons were cultured in 10 cm dishes (BioScience) for 18 days. After the culture medium was aspirated, neurons were collected with PBS (with Mg^2+^ and Ca^2+^). Collected neurons were then centrifuged for 30 seconds using a table-top centrifuge, and the pellet saved in nitrogen and stored at −20°C. Protein concentration was determined using the BCA assay (Pierce™ BCA Protein Assay Kits) according to the manufacturer’s instructions. Equal amounts of protein were mixed with a sample reducing agent (10X; Invitrogen, NuPAGE) and LDS Sample Buffer (4X; Invitrogen, NuPAGE) and denatured at 95 °C for 5 min before gel electrophoresis.

Proteins were separated by SDS–PAGE 4–15% TGX Gels (BioRad) and after transferred onto Nitrocellulose membranes (Trans-Blot Turbo, BioRad) using a semi-dry/wet transfer system. Membranes were blocked for 1 h at room temperature in Intercept TBS Blocking Buffer (Licor). Primary antibodies (Table 1.) were incubated overnight at 4 °C in blocking buffer. After washing in TBS buffer containing 0.1% Tween-20, membranes were incubated with secondary antibodies for 1h at room temperature. Protein bands were visualized using Li-Cor Odyssey DLx Acquisition Software. Band intensities were analyzed using ImageJ/Fiji.

#### Immunofluorescence confocal microscopy

Primary neuron cultures were fixed in PBS with 4% paraformaldehyde and 0.1% glutaraldehyde (Electron Microscopy Sciences), and 4% sucrose (Fisher Scientific) for 20 min at room temperature before washing 3 times with PBS. Fixed cultured hippocampal neurons were permeabilized with 0.5% Triton X-100 in PBS, pH 7.4, for 15 minutes and blocked in PBS containing 4% goat serum (Gibco) for 1 hour. Fixed neurons were incubated overnight with primary antibodies (Table S1) in PBS containing 4% goat serum (Gibco) at 4°C. PBS with 4% bovine serum albumin (Sigma) was used instead for goat anti-calnexin immunostaining. The samples were then washed three times in PBS (5 min each) before incubation at room temperature for 1 hour with appropriate secondary antibodies (Table S1). After washing three times in PBS (5 min each), neurons were briefly fixed in 4% paraformaldehyde (5 min) post-staining. All steps were performed at room temperature. The samples were stored in PBS at 4°C until imaging.

Purified synaptic vesicles from the mouse brain were a gift from J. Rubinstein lab that were purified and validated as previously reported (*43*). To visualize protein expression on purified synaptic vesicles, the purified vesicles were plated and fixed on 96-well glass-bottom plates with or without permeabilization, followed by immunofluorescence staining using primary and fluorescently labeled secondary antibodies (Table S1), as described above.

3D Confocal microscopy was performed with ZEISS LSM 980 with Airyscan 2 and image analyses of vGluT1-positive synapses, the MAP2-positive dendritic shaft, and the synapotopodin-positive spine apparatus were processed with ImageJ. Briefly, their respective markers were used to threshold the regions of interest for quantifications. Signals at the spine base were quantified using the MAP2-positive dendritic segment (approx. 2 μm) at the base of a synaptopodin-positive dendritic spine.

#### DNA PAINT-based single molecule localization microscopy

Fixed cultured hippocampal neurons were permeabilized with 0.5% Triton X-100 in PBS, pH 7.4, for 15 min and blocked in PBS containing 4% goat serum (Gibco) for 1 hour. For DNA PAINT single-molecule localization microscopy, primary antibody staining was performed overnight in PBS containing 4% goat serum (Gibco), followed by three washes with PBS for 5 min each. The samples were then incubated for 1 hour with secondary antibodies for DNA PAINT including anti-mouse antibodies conjugated to a single-stranded DNA oligo (P1 docking), and anti-rabbit antibodies conjugated to a P2-docking oligo (1:1000) (Massive Photonics). After washing three times in PBS (5 min each), neurons were briefly fixed (5 min) post-staining. Gold fiducial markers were sparsely plated via 20 minutes incubation with PBS buffer containing 20 uL gold nanoparticles stock solution (90 nm, A1190, Nanoparz). For DNA-PAINT imaging, an imaging buffer containing 1uL of P1 and 1uL of P2 imager oligo conjugated with Atto655 and Cy3B (Massive Photonics) was used. PAINT was performed on an Olympus IX83 with Abbelight SAFe Nexus, and dual-CMOS, Hamamatsu C15440-S0UP camera (Abbelight). A 100× oil-immersion objective (Olympus Immoil- F30CC; Numerical aperture 1.518) was used in combination with a motorized TIRF illuminator. Image acquisition was performed with 640nm (244 mW) and 561nm (130 mW), and the Abbelight Neo software package. Epifluorescence images of vGluT and MAP2 were collected as reference images for excitatory synapses and neuronal dendrites, as previously reported (*15*).

DNA-PAINT data was processed as previously (*15*). Briefly, the images were reconstructed with Picasso : Localize, a module of the Picasso software package (github.com/jungmannlab/picasso), by applying a minimal net gradient of 15000. Following drift-correction data using Picasso : Render, the data was filtered using Picasso : Filter. Raw localizations within a maximal distance of 6X measured localization precision and showing a maximum number of transient dark frames of 20 were linked together, resulting in a single, linked localization event. DNA PAINT-based protein copy-number analysis was performed as previously (*15*, *16*). Briefly, the localizations masked by MAP2 and vGluT1 epifluorescence signals were retained as dendritic and synaptic localizations. We calculated the average number of localization events for a single protein copy using the smallest population of clusters in the number-of-linked-localization distribution for all the identified clusters, with a localization precision (measured by the nearest-neighbor analysis) of approx. 10 nm. To estimated protein copy-number at synapses, clusters were picked within the vGluT regions-of-interest. The total number of localizations divided by the average localization number of a single protein copy resulted in an estimated copy number for the protein of interest at a synapse.

#### Immunofluorescence microscopy of mouse brain slices

Mice of either sex were sacrificed by cervical dislocation, and their brains were immediately extracted and fixed in 4% PFA overnight at 4 °C. After fixation, the samples were washed three times in 1xPBS for 10 minutes and subsequently stored in PBS until sectioning. Coronal sections were cut at a thickness of 50 µm using a Leica VT1200S vibratome (Leica Microsystems, Denmark) and stored in PBS at 4 °C until further processing for immunohistochemistry.

Coronal slices of mouse brain tissue were blocked with 0.5% Triton X-100 and 4% goat serum in PBS, hereafter referred to as blocking buffer, at room temperature for 4-6 hours. After blocking, slices were incubated with primary antibodies (Table 1) diluted in blocking buffer for 24-48 hours at 4 °C. The slices were then washed three times in PBS (15 minutes each) and incubated with anti–mouse secondary antibody conjugated with Alexa Fluor 488 (1:1000; Invitrogen), anti-rabbit antibody conjugated with Alex Fluor 568 (1:1000; Invitrogen) and anti-chicken antibody 633 (1:1000; Biotium) diluted in blocking buffer for 5 hours at room temperature or overnight at 4 °C in the dark. From this point onward, all steps were done in the dark to protect the fluorescent antibodies from photobleaching. The slices were washed twice for 10 minutes in PBS and then stained with DAPI in PBS for an additional 15 minutes before washing with PBS for 3 times. The slices were kept in PBS until mounting.

For confocal imaging, the tissue slices were transferred onto angled glass slides submerged in PBS, unfolded with a paintbrush, and positioned flat. Slides were removed, blotted, briefly air-dried, then dipped in demineralized water for 2 seconds to remove salts. After final drying, mounting medium (Agilent, S3023) was applied, and a coverslip (VWR, 631-0150) was placed without trapping air bubbles. Slides were dried completely before imaging and stored at 4 °C.

#### Expansion Microscopy

4X expansion microscopy of fixed brain slices and cultured neurons expansion was performed as previously described (*51*). Briefly, immunostained samples were incubated overnight at room temperature in a solution of AcX (Thermo Fisher, A20770) at a concentration of 10 mg/mL in DMSO (Thermo Fisher, D12345), diluted 1:100 in PBS. Following incubation, the samples were washed twice with PBS for 15 minutes each. For the slices, the samples were then incubated with Stock X (Table 2) for 1 hour at 4 °C. Afterwards, each brain slice was transferred to a MatTek glass-bottom dish (P35G-1.5-14-C) filled with a gelling solution which included: Stock X (Table 2), 4HT (Alfa Aesar, A12497) at 0.1 mg/mL, TEMED (Thermo Fisher, 17919) at 0.02 mg/mL, and APS (Thermo Fisher, 17874) at 0.02 mg/mL, in a final ratio of 47:1:1:1. Gelation was carried out for 2 hours at 37 °C before enzymatic digestion and expansion in distilled water. Similarly, cultured neurons were incubated at 37 °C for 1.5 h in the gelation solution followed by enzymatic digestion and expansion in distilled water.

#### Statistics and data visualization

All experiments have been reproduced with three biological replicates unless otherwise noted. For confocal microscopy of cultured neurons *in vitro*, experiments were repeated with at least three neuronal preparations. For DNA PAINT, at least six neurons from three neuronal preparations were imaged.

Protein localization motifs were numerically represented using a character-level k-mer approach. Motif sequences were tokenized into overlapping 2–3 amino acid n-grams using a CountVectorizer (scikit-learn), generating high-dimensional count-based embeddings for each motif. These embeddings were projected into two dimensions using Uniform Manifold Approximation and Projection (UMAP; n_neighbors=15, min_dist=0.1, cosine distance) to visualize sequence-level similarity. To quantify motif reuse across genes, each motif was assigned a gene-count value corresponding to the number of genes in which the motif occurred. Motif reusability is calculated using Shannon entropy over genes:

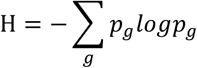

normalized by log (N_genes).

**Table S1.** Table of antibodies used in the study.

| Target | Species | Supplier | Cat. No. | Culture | Tissue | ExM | WB |
| --- | --- | --- | --- | --- | --- | --- | --- |
| Anti SERCA2 | Rb | Abcam | ab3625 | 1:500 | 1:500 | 1:500 | 1:500 |
| Anti PMCA | Ms | Invitrogen | MA3-914 | 1:500 |  |  | 1:500 |
| Anti RyR1&2 | Ms | Invitrogen | 11595802 | 1:500 |  |  | 1:500 |
| Anti vGluT1 | Ms | Synaptic Systems | 135511 | 1:500 |  |  |  |
| Anti vGluT1 | Rb | Abcam | ab227805 | 1:500 |  |  |  |
| Anti PSD-95 | Ms | Invitrogen | MA1-046 | 1:500 | 1:1000 | 1:1000 |  |
| Anti Synaptopodin | Ms | Synaptic Systems | 163011 | 1:500 |  |  |  |
| Anti MAP2 | Chk | Abcam | ab92434 | 1:5000 |  |  |  |
| Anti Calnexin | Gt | Abcam | ab219644 | 1:500 |  |  |  |
| Anti Synaptibrevin 2<br>(VAMP2) | Gp | Synaptic Systems | 104318 | 1:500 |  |  |  |
| Anti Homer1 | Rb | Synaptic Systems | 160003 | 1:500 |  |  |  |
| Anti vGluT1 | Chk | Synaptic Systems | 135316 | 1:1000 | 1:500 | 1:500 |  |
| Anti-synaptophysin | Rb | Abcam | Ab32127 | 1:500 |  |  |  |
| Anti rabbit secondary antibody, AlexaFluor 568 | Gt | Invitrogen | A11011 | 1:1000 |  | 1:500 |  |
| Anti mouse secondary antibody, AlexaFluor 488 | Gt | Invitrogen | A11001 | 1:1000 |  | 1:500 |  |
| Anti-chicken secondary antibody, AlexaFluor 647 | Gt | Invitrogen | A21449 | 1:1000 |  |  |  |
| Anti guinea pig secondary antibody, AlexaFluor 647 | Gt | Invitrogen | A21450 | 1:1000 |  |  |  |
| Anti Goat secondary antibody, AlexaFluor 647 | Dk | Thermo Fisher | A21447 | 1:1000 |  |  |  |
| Anti-chicken secondary antibody, 633nm | Gt | Biotium | 20126 |  |  | 1:500 |  |
| Mouse IgG IRDye 680RD | Dk | Li-cor | 926-68072 |  |  |  | 1:2000 |
| Rabbit IgG IRDye 800CW | Dk | Li-cor | 926-32213 |  |  |  | 1:2000 |

**Table S2.** Stock X for 4X Expansion Microscopy.

| Reagent | Stock solution concentration (g/100 ml) | Amount (mL) | Supplier | Cat. No. |
| --- | --- | --- | --- | --- |
| Sodium acrylate | 38 | 2.25 | Sigma | 408220 |
| Acrylamide | 50 | 0.5 | Sigma | A9099 |
| N,N- methylenebisacrylamide | 2 | 0.75 | Sigma | 146072 |
| NaCl (5M) | 29.2 | 4 | Sigma | S9625 |
| PBS (10X) | NA | 1 | NA |  |
| Water | NA | 0.9 | NA |  |

**Fig. S1.**
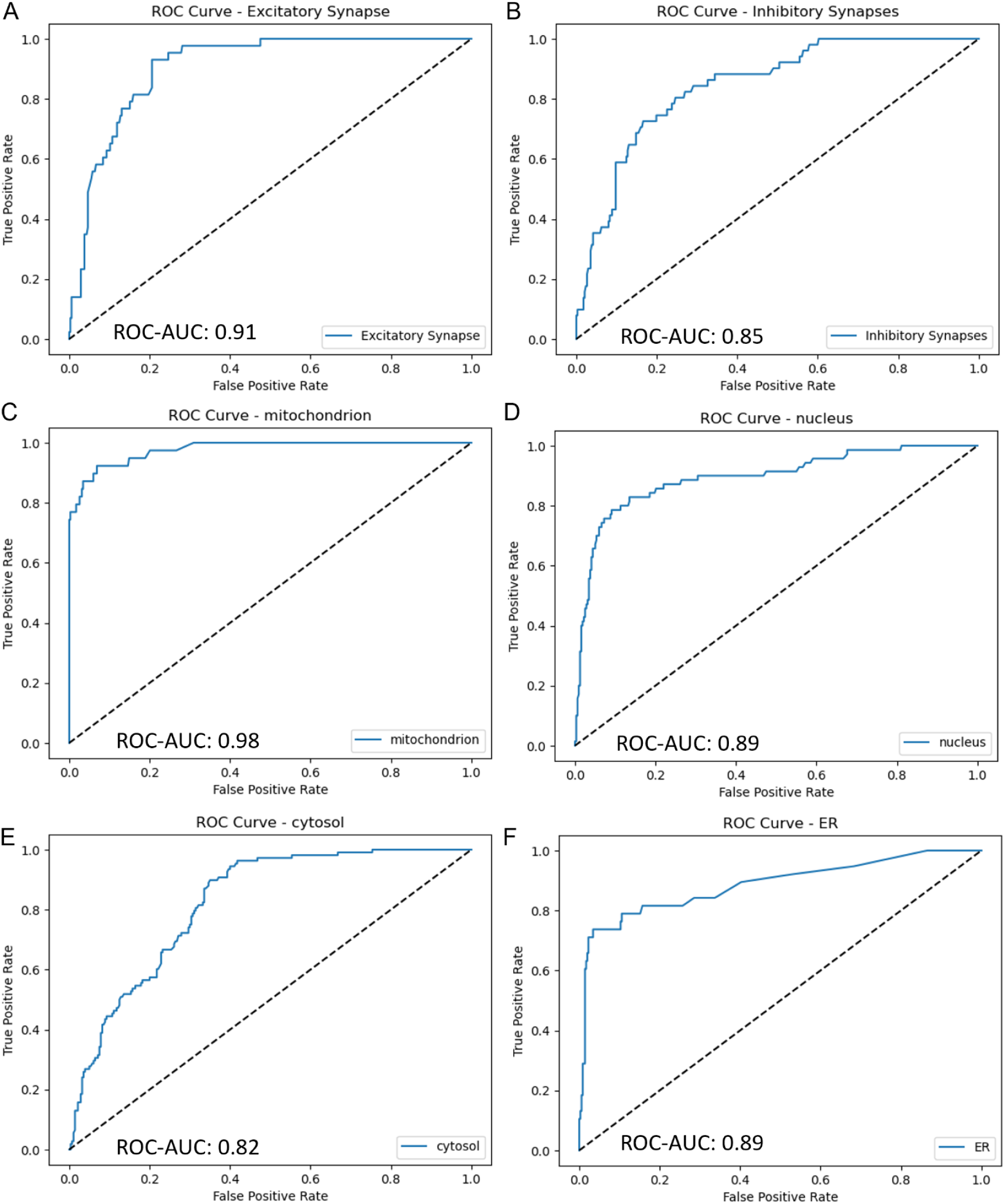
ROC-AUC curves for ESM-2-based SyGi prediction of genes at different subcellular compartments. (A-F) ROC-AUC curves for the excitatory synapse (A), the inhibitory synapse (B), the mitochondrion (C), the nucleus (D), the cytosol (E), and the ER (F).

**Fig. S2.**
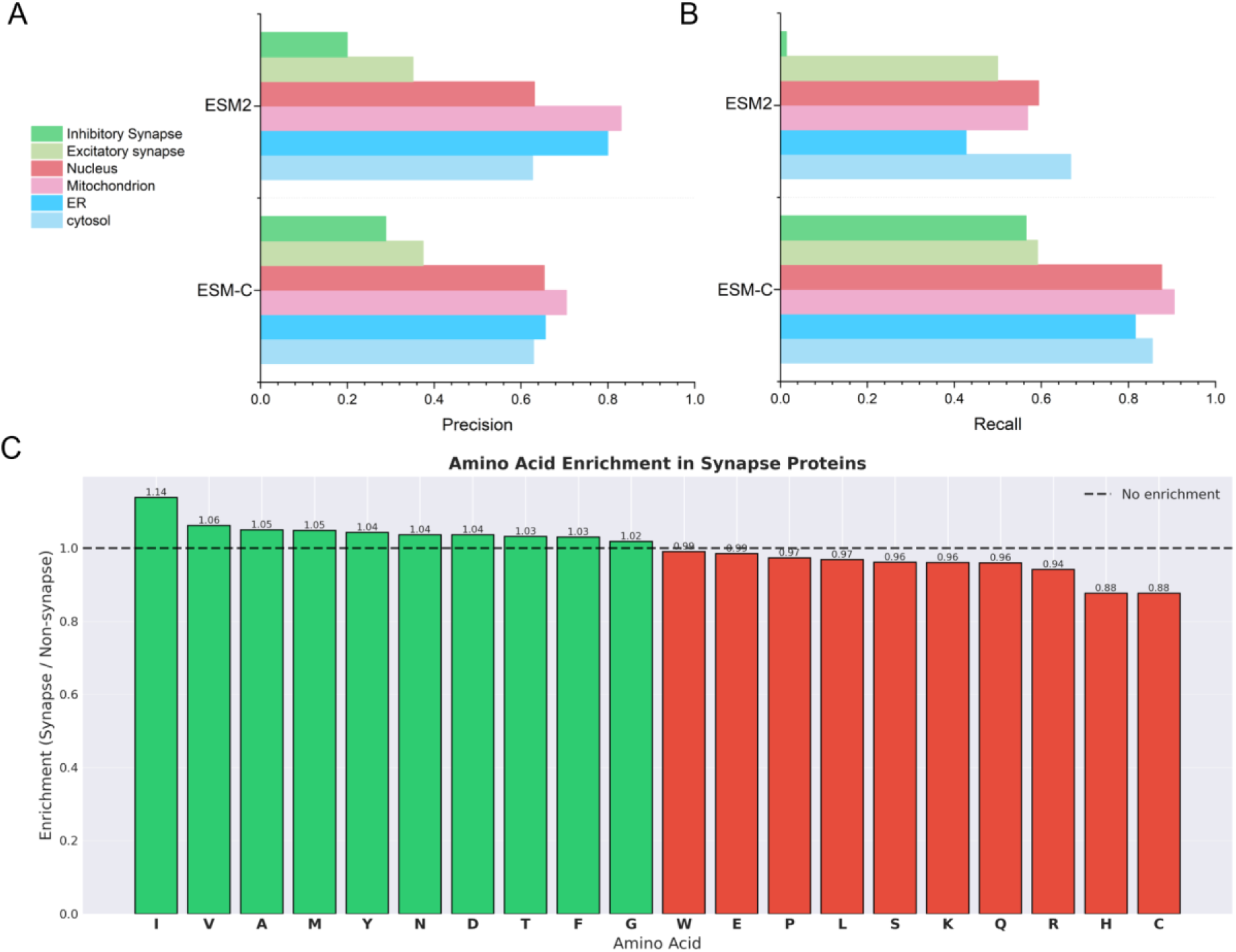
Precision and Recall performances for ESM-2- versus ESM-C-based SyGi. (A) Bar graph showing an overall better precision for ESM-2-based SyGi compared to ESM-C-based SyGi using an arbitrary confidence-score cutoff of 0.50, where Precision = True positive / (True positive + False positive). (B) Bar graph showing an overall better recall for ESM-C-based SyGi compared to ESM-2-based SyGi using an arbitrary confidence-score cutoff of 0.50, where Recall = True positive / (True positive + False negative). (C) Bar graph showing the enrichment (green) or depletion (red) of specific amino acids in synaptic protein sequences from ESM-2-based SyGi.

**Fig. S3.**
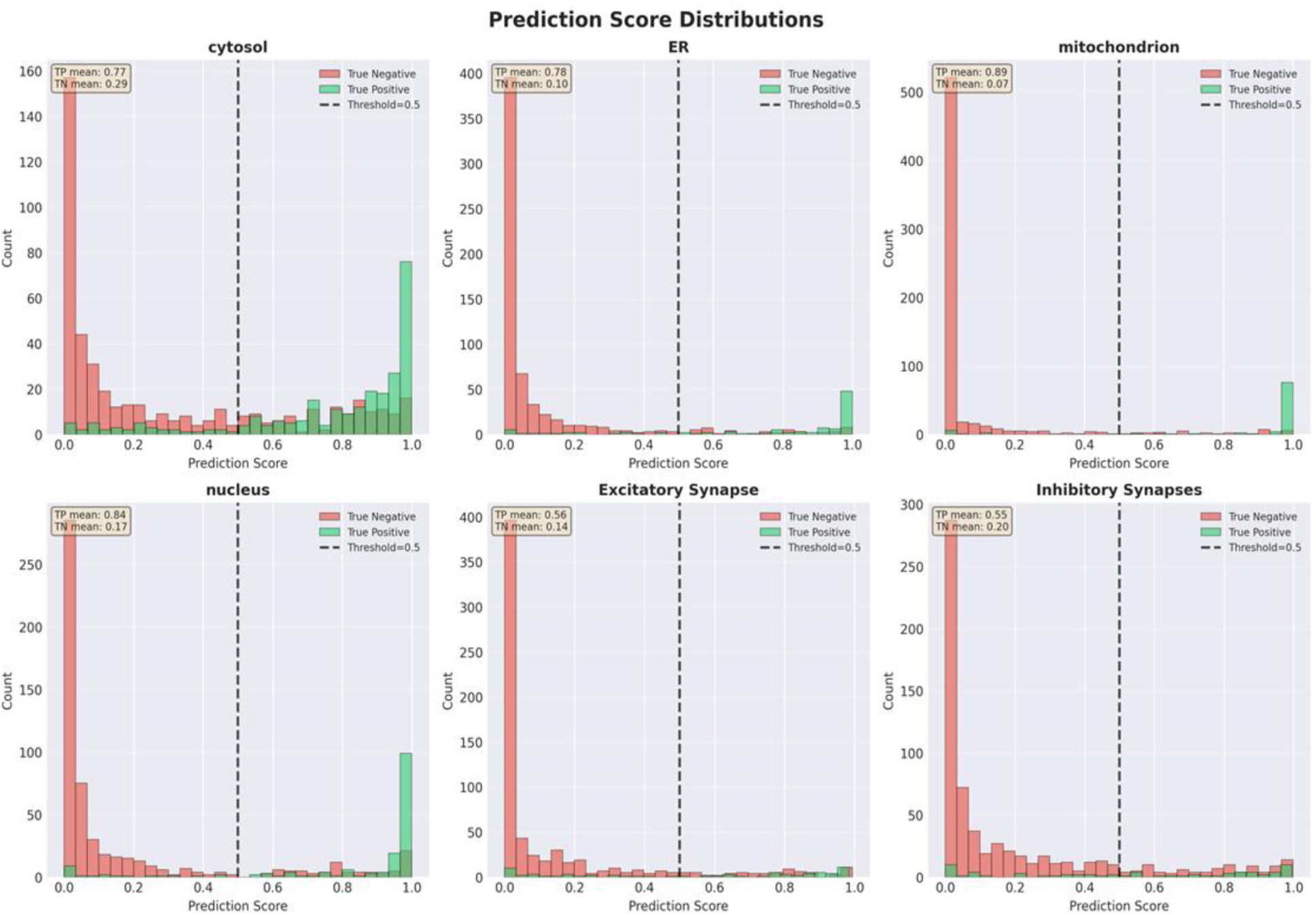
Histograms showing the gene-count distribution across the predictions scores from ESM-C based SyGi for six subcellular compartments. Dashed line indicates an arbitrary cut-off used in Figs.1, S2, & S4.

**Fig. S4.**
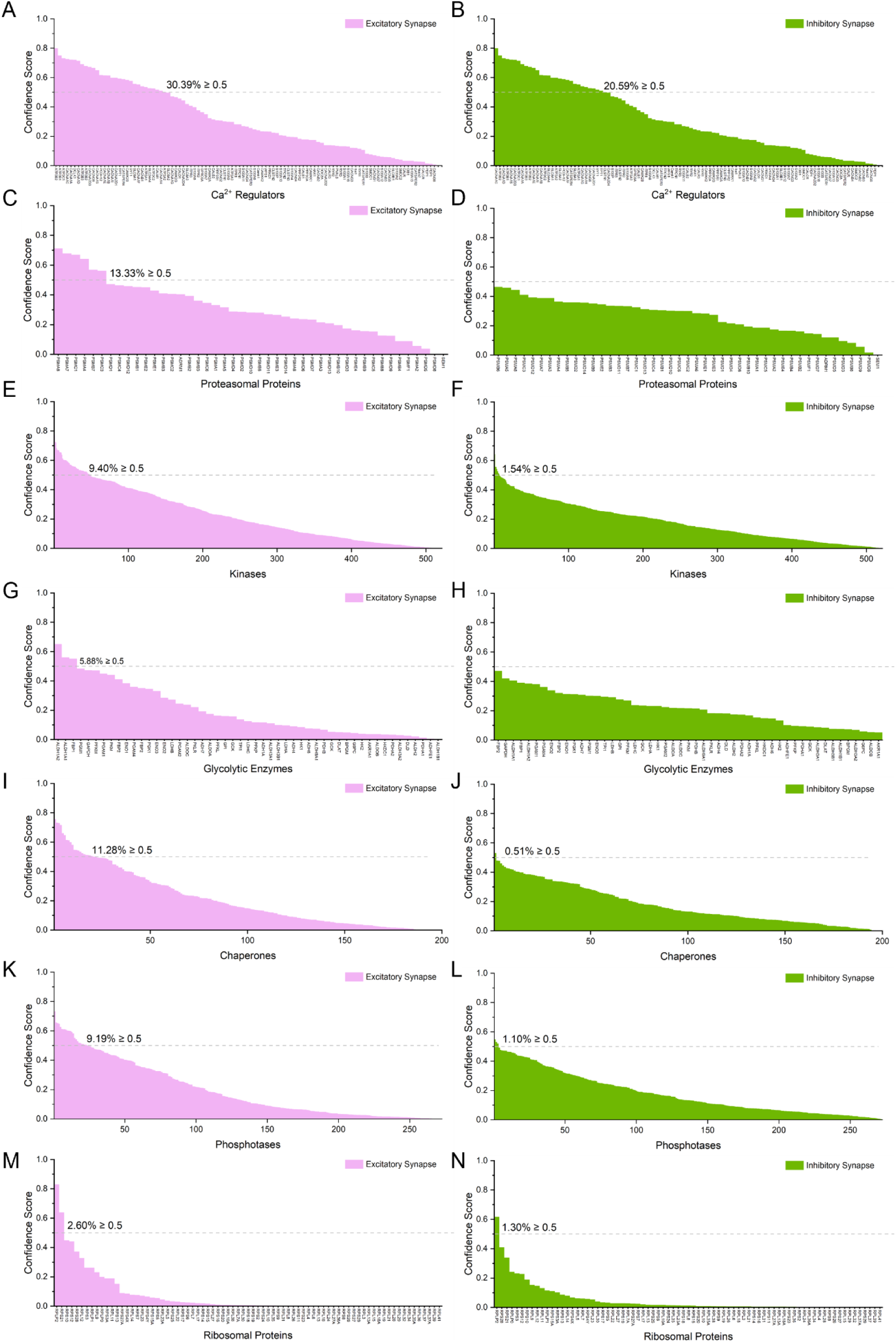
Distributions of confidence scores for synaptic localization among protein functional families for key cell biological pathways using ESM2-based SyGi. (A-N) Histogram distributions of SyGi-predicted confidence scores for calcium regulators (A-B), proteasomal proteins (C-D), cellular kinases (E-F), glycolytic enzymes (G-H), chaperones (I-J), phosphatases (K-L), ribosomal proteins (M-N). Magenta indicates score distributions for predicted excitatory synapse localization; green indicates score distributions for predicted inhibitory synapse localization. Dashed lines indicate an arbitrary cut-off for selecting high-confidence predictions.

**Fig. S5.**
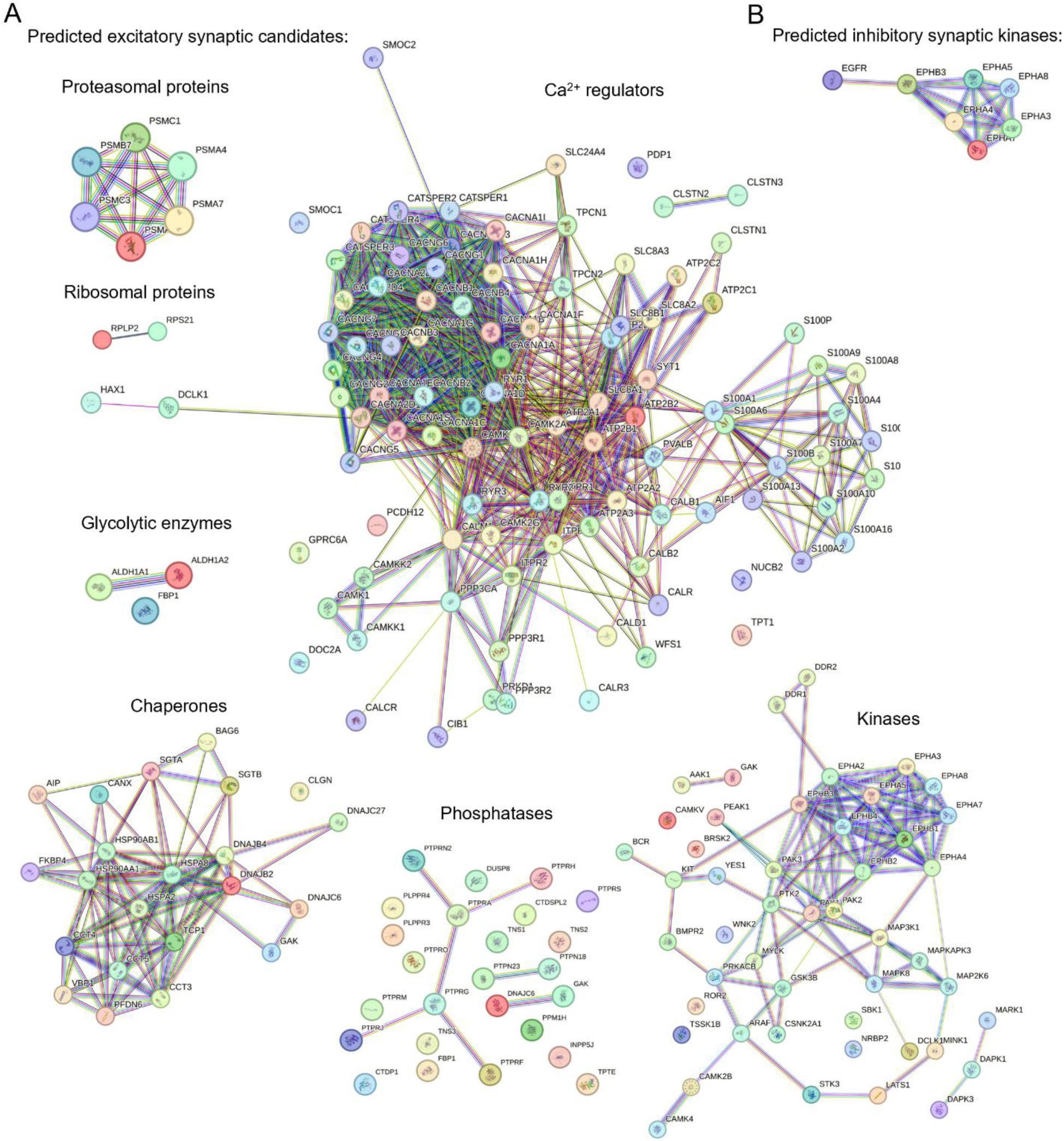
SyGi shortlists hundreds of proteins from major cellular pathways as potential synaptic components. (A) String networks of SyGi-predicted candidate genes for excitatory synaptic localization. (B) String network of SyGi-predicted candidate kinases for inhibitory synaptic localization.

**Fig. S6.**
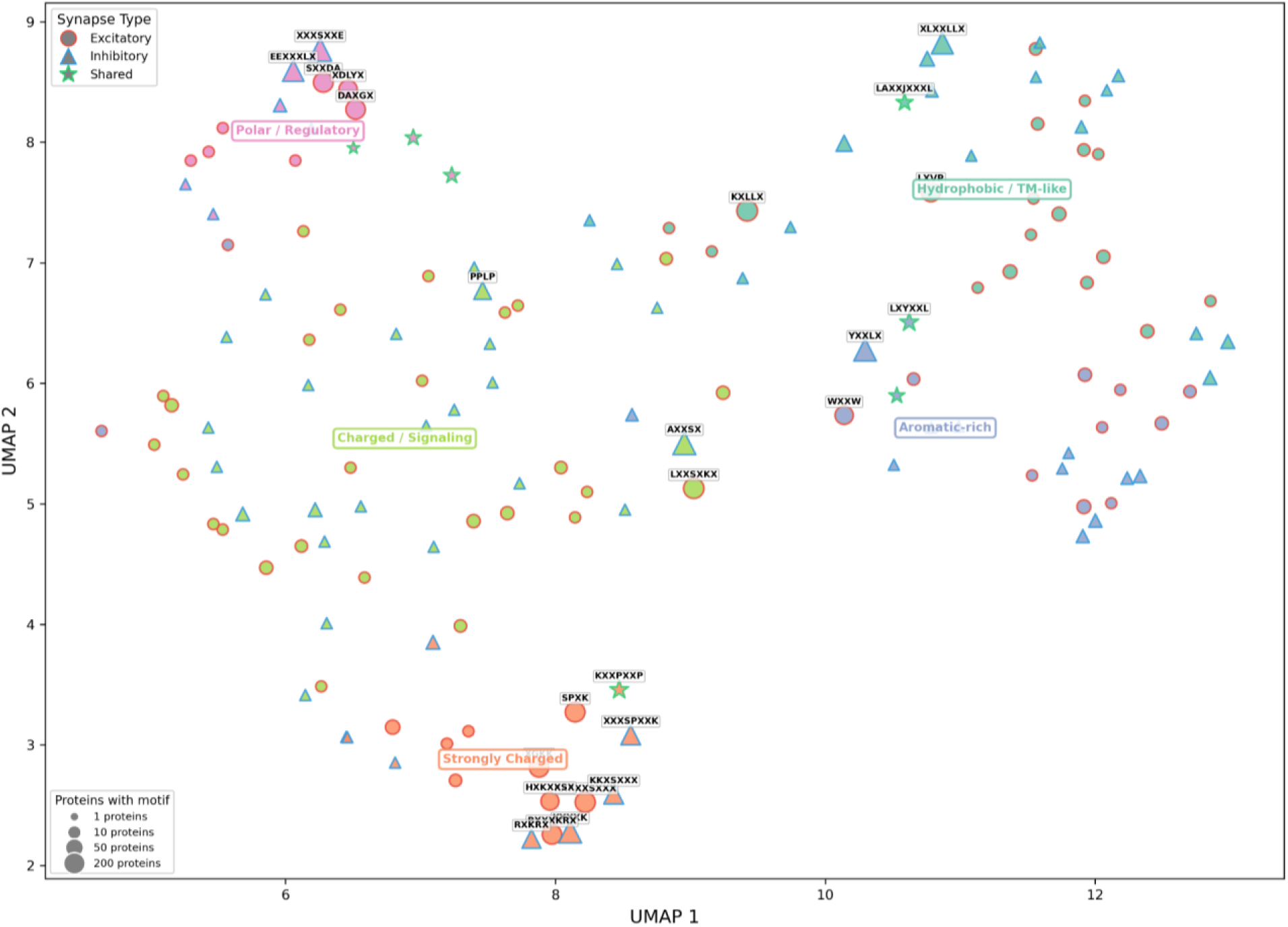
UMAP plot showing the biophysical property landscape of amino-acid motifs enriched in proteins of excitatory and inhibitory synapses (see also Supplemental Data S3).

**Fig. S7.**
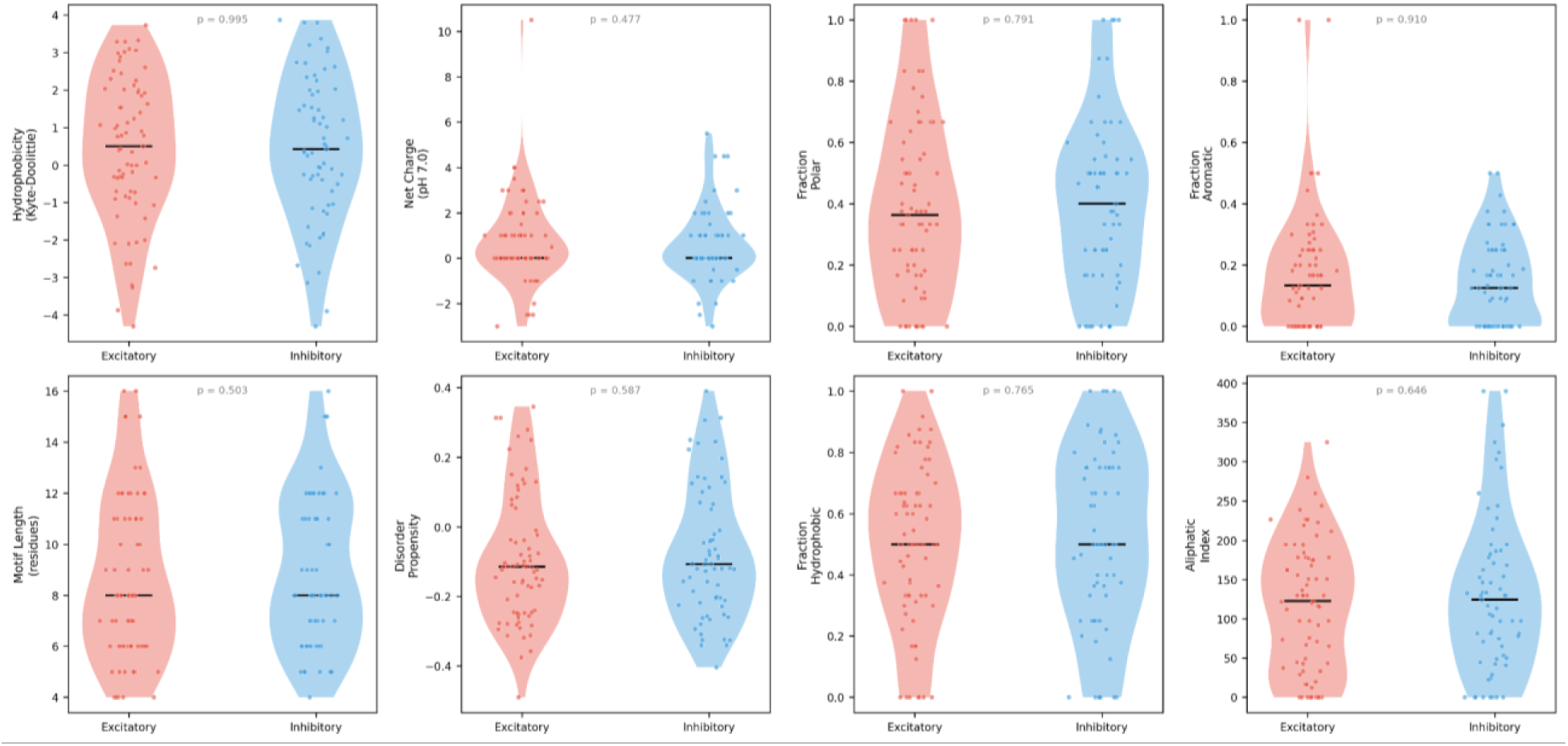
Violin plots showing a lack of significant differences between the biophysical properties of excitatory synaptic motifs and those of inhibitory synaptic motifs.

**Fig. S8.**
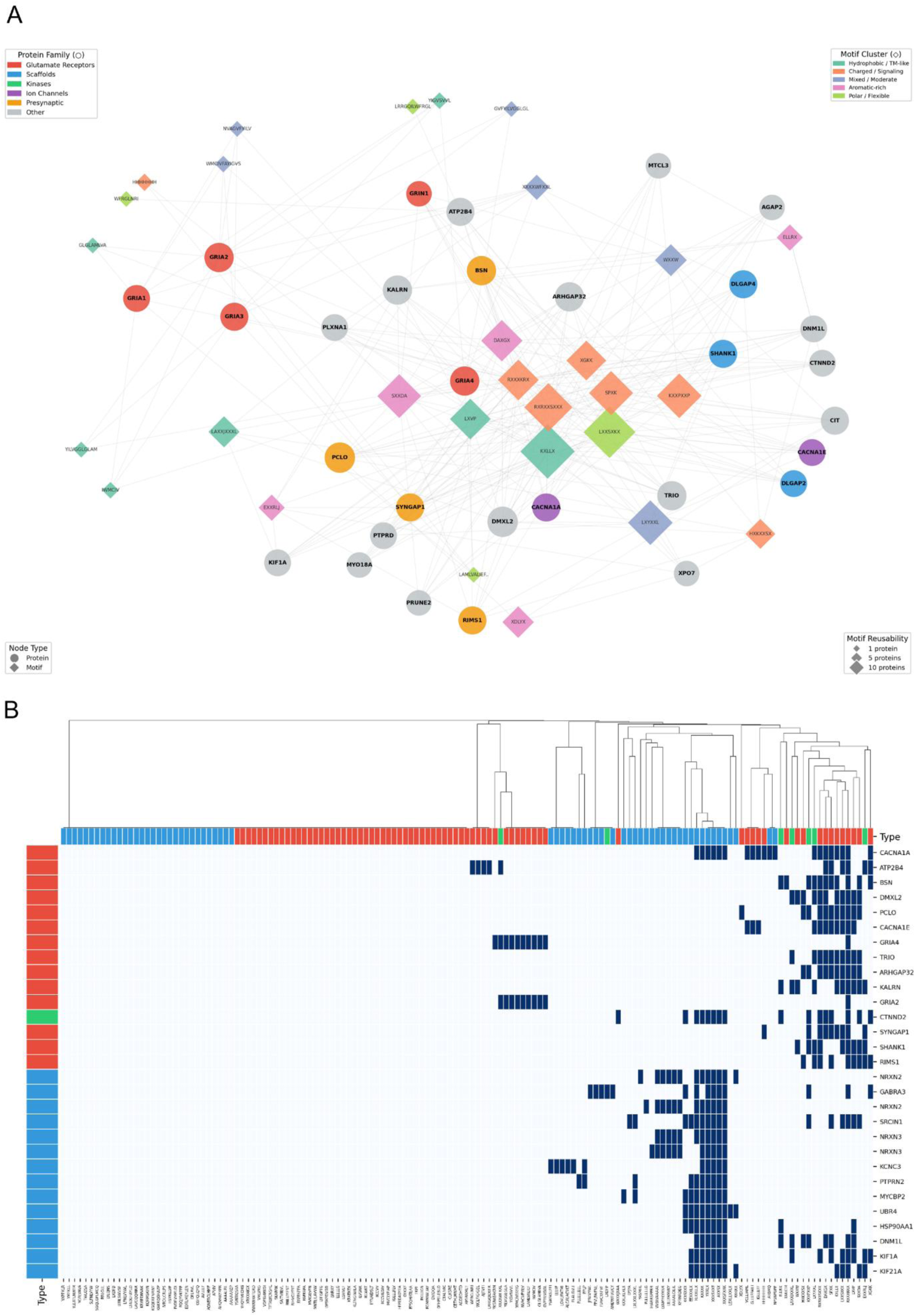
Overview of synaptic motifs shared among multiple synaptic proteins. **(A)** Bipartite network connecting excitatory synapse proteins to their short linear motifs. Circles represent synaptic proteins, colored by functional family (legend, upper left). Diamonds represent sequence motifs, colored by biophysical cluster based on hydrophobicity, charge, and aromatic content (legend, upper right). Diamond size denotes the number of proteins in this dataset containing that motif (legend, lower right). Grey lines connect proteins to motifs found within their sequences. Highly connected motifs (larger diamonds) represent recurring sequence patterns across multiple synaptic proteins, suggesting conserved functional roles. (B) Heatmap showing shared amino-acid motifs (columns) of top excitatory and inhibitory hub proteins (rows). Red denotes excitatory synapse-specific. Blue denotes inhibitory synapse-specific. Green denotes shared motifs and proteins between excitatory and inhibitory synapses.

**Fig. S9.**
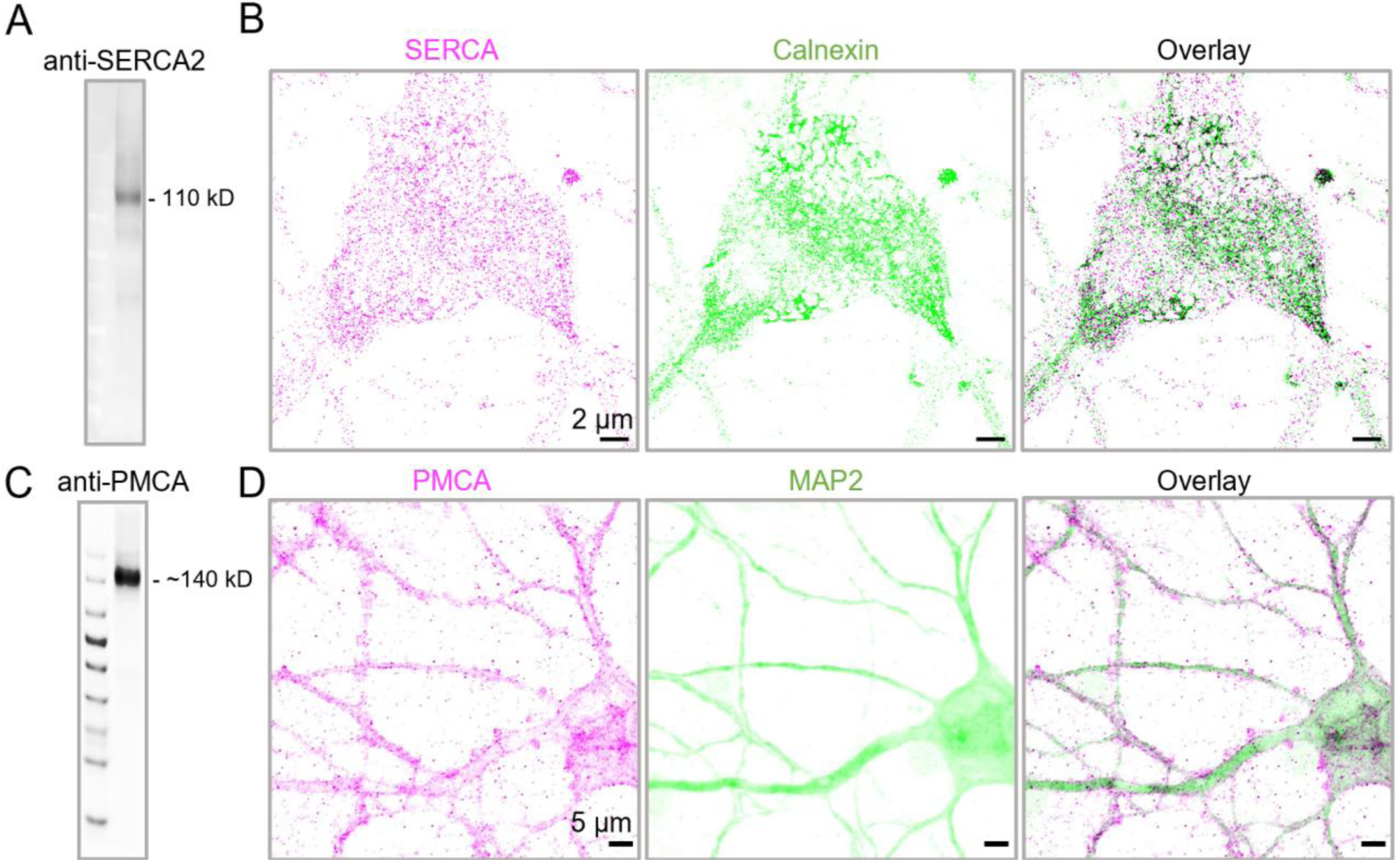
Western blotting and immunostaining validation for anti-SERCA and anti-PMCA antibodies. (A) Western blotting of anti-SERCA2 antibodies using neuronal lysates, showing the band of SERCA2 at the expected molecular mass, ∼110 kD. (B) Confocal micrograph showing the co-localization (black) of SERCA (magenta) and calnexin (green; ER marker) at the cell soma in primary neuron cultures. Scale bars: 2 μm. (C) Western blotting of anti-PMCA antibodies using neuronal lysates, showing the band of PMCA at the expected molecular mass, ∼140 kD. (D) Confocal micrograph showing the plasma-membrane expression of PMCA (magenta) in primary neurons (green). Scale bars: 5 μm.

**Fig. S10.**
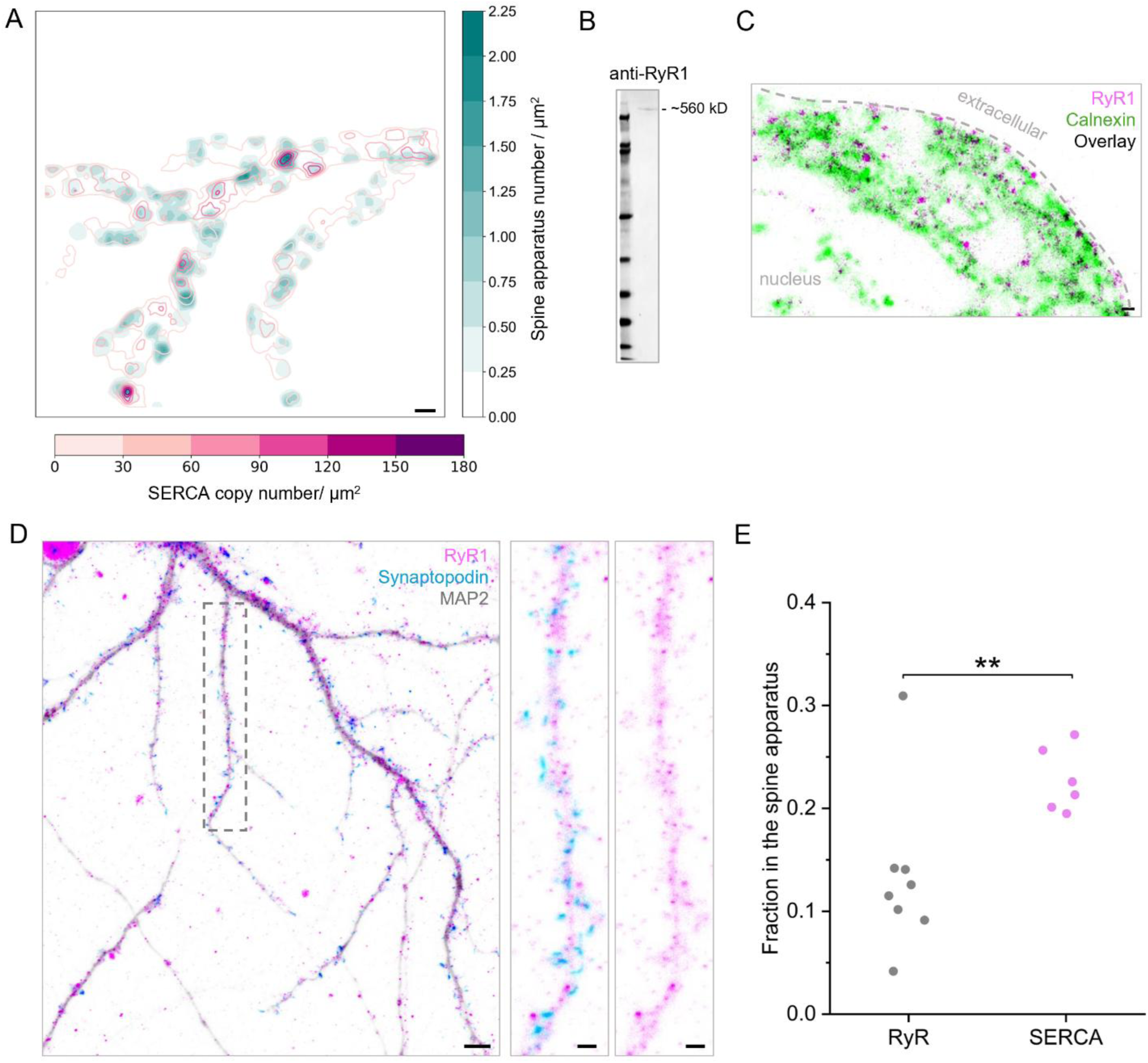
The ryanodine receptor (RyR) does not cluster at the spine apparatus. (A) Overlay between the heatmap of the spine apparatus distribution and the contour map of detected SERCA copy distribution in neuronal dendrites of Fig. 4E, showing clustering of SERCA (dark magenta contours) at the hotspots of the spine apparatus (dark teal). (B&C) Validation of anti-RyR antibodies via western blotting using mouse cortical lysates (∼560 kD band in B) and immunofluorescence confocal microscopy of ER (green in B; anti-calnexin) and RyR (magenta in C) in cultured neurons, showing the localization of RyR on ER in the cell soma (overlay in black). A zoomed-in view was shown to reveal the diffraction-limited details of ER structure. Scale bar: 1 μm. (D) Confocal micrograph showing the dendritic expression of RyR (magenta) and synaptopodin (cyan; marker for the spine apparatus) along neuronal dendrites (grey; Map2), with the overlap between RyR and synaptopodin in blue. Scale bar: 5 μm. Right inset reveals an absence of RyR clustering at the synaptopodin-positive area. Scale bars: 2 μm. (E) Column scatter plot showing a significantly higher fraction of dendritic SERCA in the synaptopodin-positive area in comparison to dendritic RyR (>20 dendritic branches of ≥ 6 neuron replicates from 3 neuronal preparations per group; two-sample T-test; p**<0.01).

**Fig. S11.**
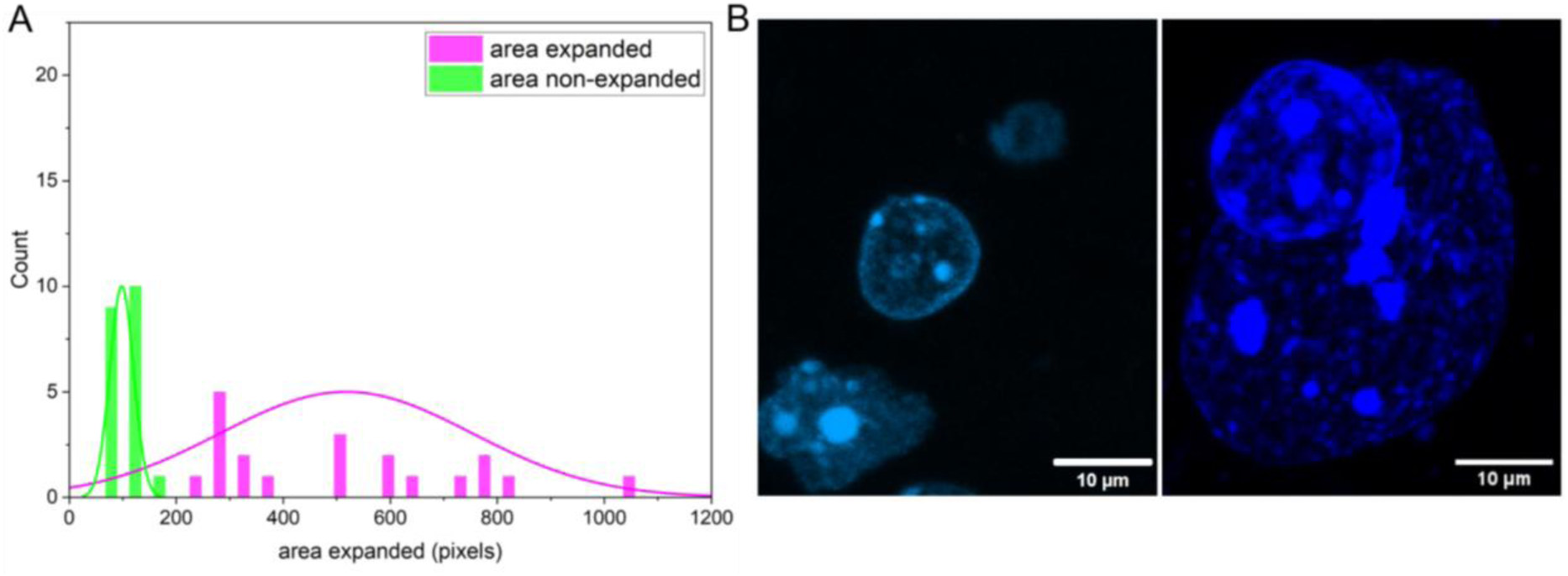
Validation for ∼4X expansion microscopy of fixed brain slices. (A) Histogram showing the size distribution of the nuclei before (green) and after (magenta) expansion in fixed brain slices. (B) Confocal micrograph showing exemplar nuclei stained by DAPI in fixed brain slices before (left) and after (right) 4X expansion. Scale bars: 10 μm.

**Fig. S12.**
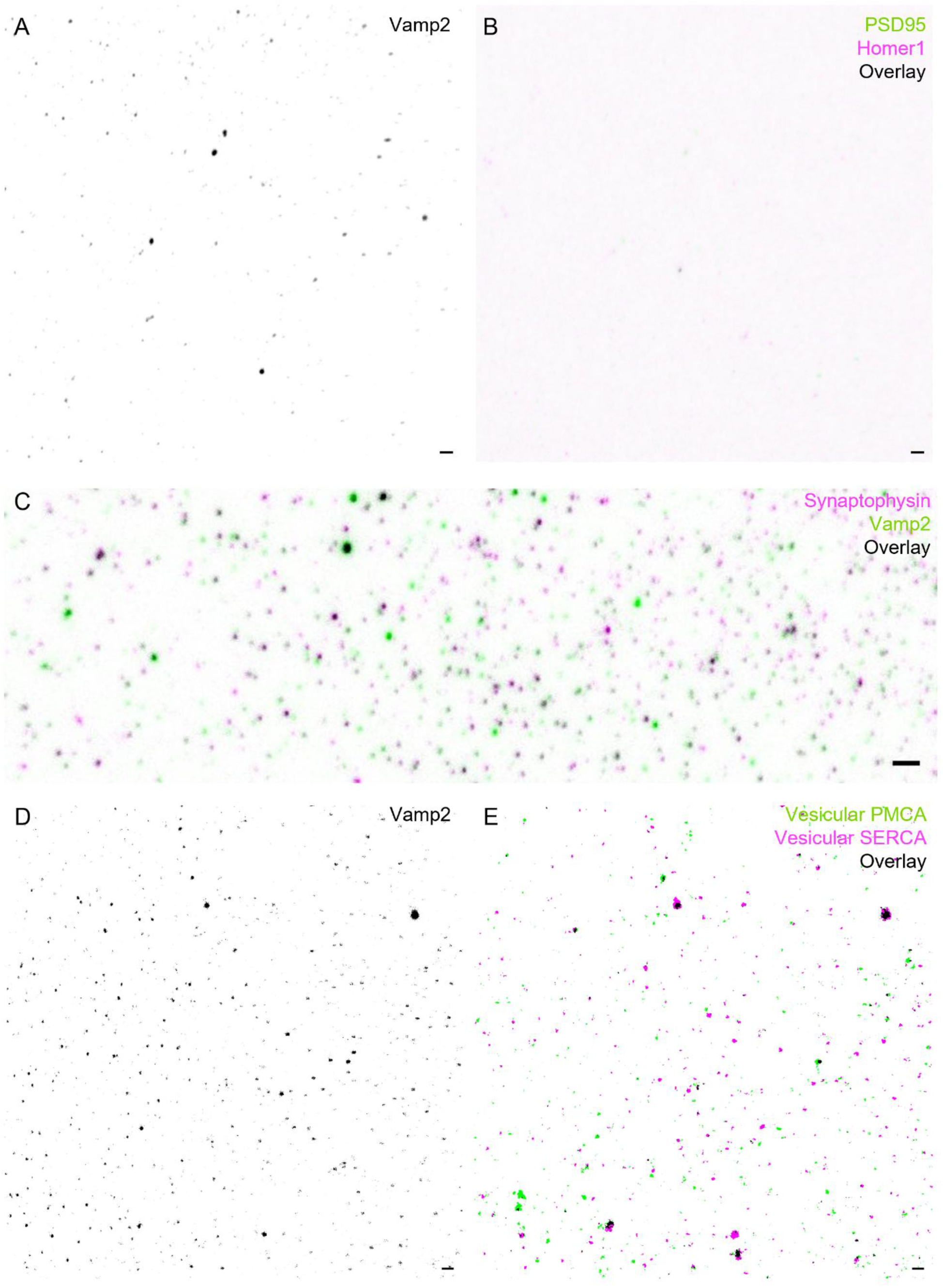
Validation for immunofluorescence microscopy of purified synaptic vesicles. (A&B) Confocal micrograph showing the absence of non-vesicular synaptic proteins PSD95 (green; B) or Homer1 (magenta; B) in Vamp2-positive puncta (A) using purified synaptic vesicles plated on glass coverslips. Scale bars: 2 μm. (C) Confocal micrograph showing the colocalization (black) of Vamp2 (green) and another vesicular protein synaptophysin (magenta) using synaptic vesicles plated on glass, with 98.4±2.7% Vamp2-positive puncta containing synaptophysin signals (averaged across 6 fields-of-view). Scale bar: 2 μm. (D&E) Confocal micrographs showing the presence of vesicular SERCA (magenta; E) and PMCA (green; E) in Vamp2-masked areas (D) using purified synaptic vesicles fixed and immunostained without permeabilization. Scale bar: 2 μm.

### Data S1. (separate file)

Curated genes and subcellular annotations for SyGi. This file contains the tabulated data behind Fig. 1C.

### Data S2. (separate file)

Gene list of cellular pathways and their prediction scores by SyGi. This file contains the tabulated data behind Fig. 1H.

### Data S3. (separate file)

Motif properties. This file contains tabulated data behind Fig. 2.

